# A Single Mechanism of Biogenesis, Initiated and Directed by PIWI Proteins, Explains piRNA Production in Most Animals

**DOI:** 10.1101/261545

**Authors:** Ildar Gainetdinov, Cansu Colpan, Amena Arif, Katharine Cecchini, Phillip D. Zamore

**Author notes:** These authors contributed equally (Ildar Gainetdinov, Cansu Colpan). Correspondence (PDZ).

## Abstract

In animals, piRNAs guide PIWI-proteins to silence transposons and regulate gene expression. The mechanisms for making piRNAs have been proposed to differ among cell types, tissues, and animals. Our data instead suggest a single model that explains piRNA production in most animals. piRNAs initiate piRNA production by guiding PIWI proteins to slice precursor transcripts. Next, PIWI proteins direct the stepwise fragmentation of the sliced precursor transcripts, yielding tail-to-head strings of phased pre-piRNAs. Our analyses detect evidence for this piRNA biogenesis strategy across an evolutionarily broad range of animals including humans. Thus, PIWI proteins initiate and sustain piRNA biogenesis by the same mechanism in species whose last common ancestor predates the branching of most animal lineages. The unified model places PIWI-clade Argonautes at the center of piRNA biology and suggests that the ancestral animal—the Urmetazoan—used PIWI proteins both to generate piRNA guides and to execute piRNA function.

## INTRODUCTION

Uniquely, animals produce PIWI-interacting RNAs (piRNAs), a special class of small RNAs that guides PIWI proteins to silence transposons and regulate gene expression (Girard et al., 2006; Aravin et al., 2006; Vagin et al., 2006; Saito et al., 2006; Grivna et al., 2006; Houwing et al., 2007; Batista et al., 2008; Das et al., 2008). piRNAs complementary to transposons ensure genomic stability in the male or female germline in animals as diverse as scorpions, honey bees, and mice; in many arthropods, piRNAs also protect somatic tissues from both transposons and RNA viruses (Siomi et al., 2011; Czech and Hannon, 2016; Morazzani et al., 2012; Miesen et al., 2015; Vodovar et al., 2012; Lewis et al., 2017). Remarkably, in silk moth oocytes, a single piRNA helps determine sex (Kiuchi et al., 2014). In the testes of mammals, distinct classes of piRNAs (1) direct DNA and histone methylation to transposon sequences during embryogenesis (Aravin et al., 2008; Kuramochi-Miyagawa et al., 2008; Pezic et al., 2014), (2) post-transcriptionally repress transposons later in spermatogenesis (Reuter et al., 2011; Di Giacomo et al., 2013; Inoue et al., 2017), and (3) ensure completion of meiosis and successful spermiogenesis (Aravin et al., 2006; Girard et al., 2006; Grivna et al., 2006; Vourekas et al., 2012; Watanabe et al., 2014; Gou et al., 2014; Goh et al., 2015; Zhang et al., 2015).

Studies of oogenesis in flies and spermatogenesis in mice suggest that germline piRNA biogenesis can be divided into primary and secondary pathways. Primary piRNAs (Figure 1, at left) are generated from long, single-stranded RNAs that are transcribed from genomic loci called piRNA clusters (Brennecke et al., 2007; Li et al., 2013). These long precursor transcripts are fragmented by endonucleolytic cleavage—hypothesized to be catalyzed by the mitochondrial protein Zucchini/PLD6—producing tail-to-head, phased precursor piRNAs (pre-piRNAs; Figure 1, at left; Ipsaro et al., 2012; Nishimasu et al., 2012; Mohn et al., 2015; Han et al., 2015; Ding et al., 2017). Each pre-piRNA begins with a 5′ monophosphate, a prerequisite for loading RNA into nearly all Argonaute proteins (Nykanen et al., 2001; Ma et al., 2005; Parker et al., 2005; Wang et al., 2009; Frank et al., 2010; Boland et al., 2011; Kawaoka et al., 2011; Elkayam et al., 2012; Schirle and MacRae, 2012; Schirle et al., 2014; Cora et al., 2014; Wang et al., 2014; Matsumoto et al., 2016). Once bound to a PIWI protein, the 3′ ends of pre-piRNAs are trimmed by the single-stranded-RNA exonuclease, Trimmer/PNLDC1, to a length characteristic of the receiving PIWI protein (Kawaoka et al., 2011; Tang et al., 2016; Izumi et al., 2016; Ding et al., 2017; Zhang et al., 2017; Nishimura et al., 2018). (Flies lack a Trimmer/PNLDC1 homolog, and instead piRNA 3′ ends are resected by the miRNA-trimming enzyme Nibbler; [Han et al., 2011; Liu et al., 2011; Feltzin et al., 2015; Hayashi et al., 2016].) Finally, the small RNA methyltransferase Hen1/HENMT1 adds a 2′-*O*-methyl moiety to the 3′ ends of the mature piRNAs (Kirino and Mourelatos, 2007b; Horwich et al., 2007; Saito et al., 2007; Kirino and Mourelatos, 2007a; Ohara et al., 2007; Lim et al., 2015).

**Figure 1.**
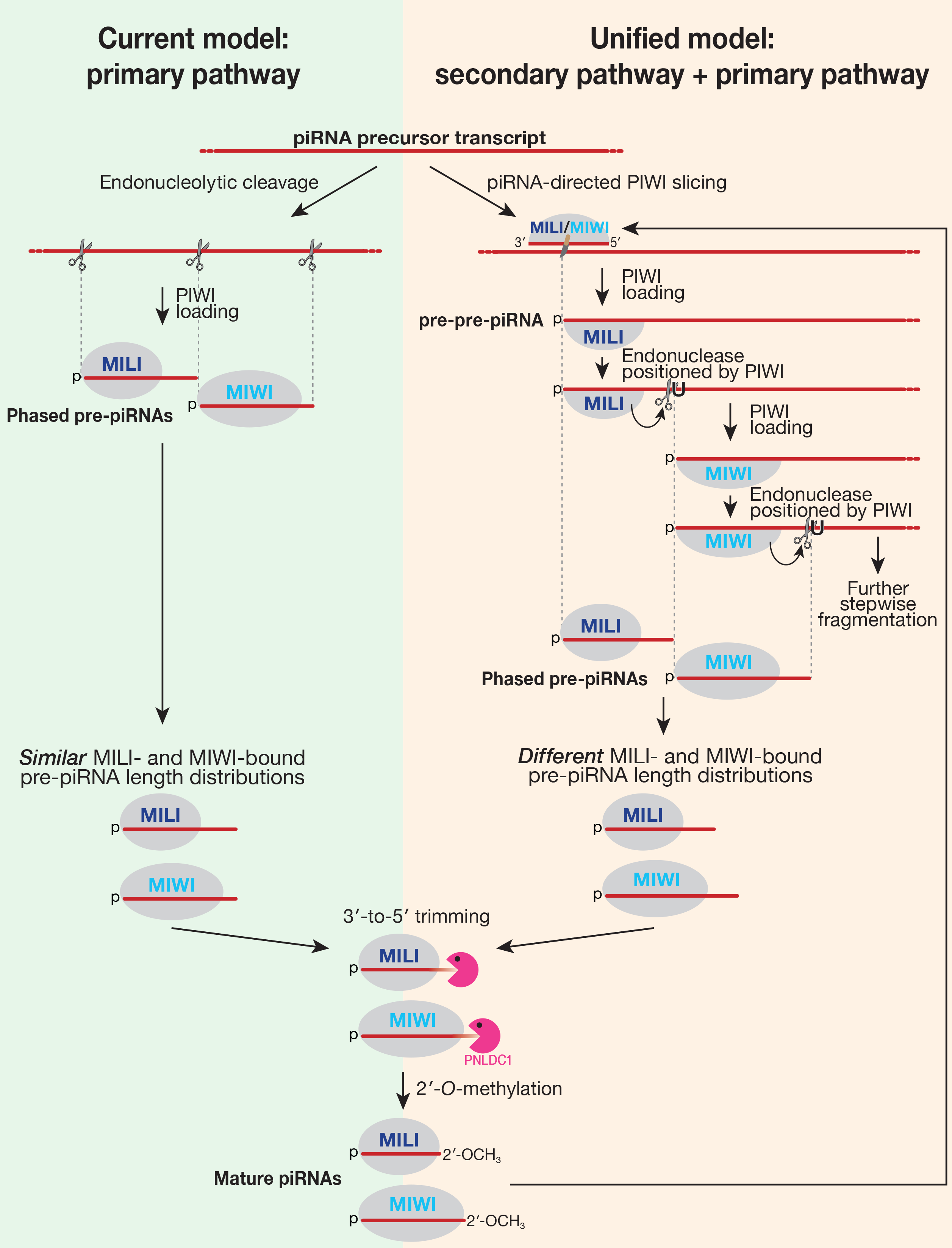
The Current Model for Primary piRNA Biogenesis in Post-Natal Mouse Testis and the Unified Model for piRNA Biogenesis in Animals.

In the secondary piRNA biogenesis pathway or “ping-pong” cycle (Brennecke et al., 2007; Gunawardane et al., 2007), piRNA-directed, PIWI-catalyzed slicing of a target transcript creates an RNA fragment bearing a 5′ monophosphate (Wang et al., 2014). This RNA fragment acts as a pre-piRNA precursor (pre-pre-piRNA), binds to a PIWI protein, and ultimately generates a new secondary piRNA with 10 nt complementarity to the piRNA that produced it.

According to the current model, distinct mechanisms produce piRNAs in different cell types and tissues and in different animals (Reuter et al., 2011; Beyret et al., 2012; Vourekas et al., 2012; Nishida et al., 2018). For example, fly oocytes and mouse embryonic male germ cells produce both primary and secondary piRNAs, and the secondary pathway counterintuitively initiates biogenesis of primary piRNAs: piRNA-directed, PIWI-catalyzed slicing of a long precursor transcript creates a secondary piRNA followed by a series of phased pre-piRNAs, which then mature into primary piRNAs (Mohn et al., 2015; Han et al., 2015; Senti et al., 2015; Wang et al., 2015; Yang et al., 2016). In contrast, the current model for mouse pachytene piRNA biogenesis proposes that “piRNA biogenesis in post-natal male germ cells differs strikingly from that in embryonic [germ] cells, because the majority of piRNAs are produced only by primary biogenesis after birth” (Nishimura et al., 2018; Figure 1, at left). What initiates primary piRNA production in the post-natal mammalian testis is unknown. Finally, recent work on cultured silk moth BmN4 cells reported that “*Bombyx* produces no phased piRNAs” (Nishida et al., 2018), implying that piRNAs in the silk moth are produced solely through the secondary pathway, i.e., by ping-pong amplification.

Here, we report that a single mechanism can explain piRNA biogenesis in mammals, insects, and likely all other animal germ cells (Figure 1, at right). We show that the secondary piRNA pathway—piRNA-guided slicing by a PIWI protein—initiates primary piRNA biogenesis in the post-natal mouse testis. Thus, the same fundamental strategy explains how piRNA production begins in the female germline in flies and the pre- and post-natal male germline in mammals. Moreover, both in mice and in flies, PIWI proteins directly participate in primary piRNA production by directing the stepwise fragmentation of long piRNA precursor transcripts: the site of endonucleolytic cleavage that simultaneously establishes the 3′ end of a pre-piRNA and the 5′ end of the next primary piRNA is determined by the PIWI protein bound to the precursor transcript’s 5′ end. Analysis of piRNA sequences detects evidence for this same piRNA biogenesis strategy across an evolutionary broad range of animals. Thus, a common biogenesis architecture describes how PIWI proteins initiate and sustain production of their own piRNA guides in animals separated by almost a billion years of evolution (Hedges et al., 2015; Kumar et al., 2017). We hypothesize that the piRNA pathway in the ancestral animal consisted only of PIWI-clade Argonautes that both generated their own piRNA guides and executed piRNA function.

## RESULTS

### Phased Pre-piRNAs are a General Feature of Primary piRNA Biogenesis in Bilateral Animals

In the dipteran insect *Drosophila melanogaster* (fruit fly) and the mammal *Mus musculus* (house mouse), primary piRNA biogenesis pathway produces tail-to-head strings of phased pre-piRNAs, which are further 3′-to-5′ trimmed to yield mature primary piRNAs (Mohn et al., 2015; Han et al., 2015; Ding et al., 2017). We asked whether this strategy for making piRNAs was broadly conserved, analyzing piRNA sequencing data from 33 non-model species spanning ~950 million years of evolution (Figure 2). In mice and flies, genetic mutation or RNAi depletion of *papi*/*Tdrkh*, *nibbler*, or *Trimmer*/*Pnldc1* blocks piRNA maturation, allowing the detection of tail-to-head strings of untrimmed pre-piRNAs, the hallmark of the phased, primary piRNA pathway (Saxe et al., 2013; Mohn et al., 2015; Han et al., 2015; Hayashi et al., 2016; Ding et al., 2017). Such loss-of-function strategies are typically not available for non-model species. However, studies in flies have shown that phasing can be detected among mature piRNAs when the extent of pre-piRNA trimming is small, i.e., when untrimmed pre-piRNAs and trimmed mature piRNAs are similar in length (Mohn et al., 2015; Han et al., 2015).

**Figure 2.**
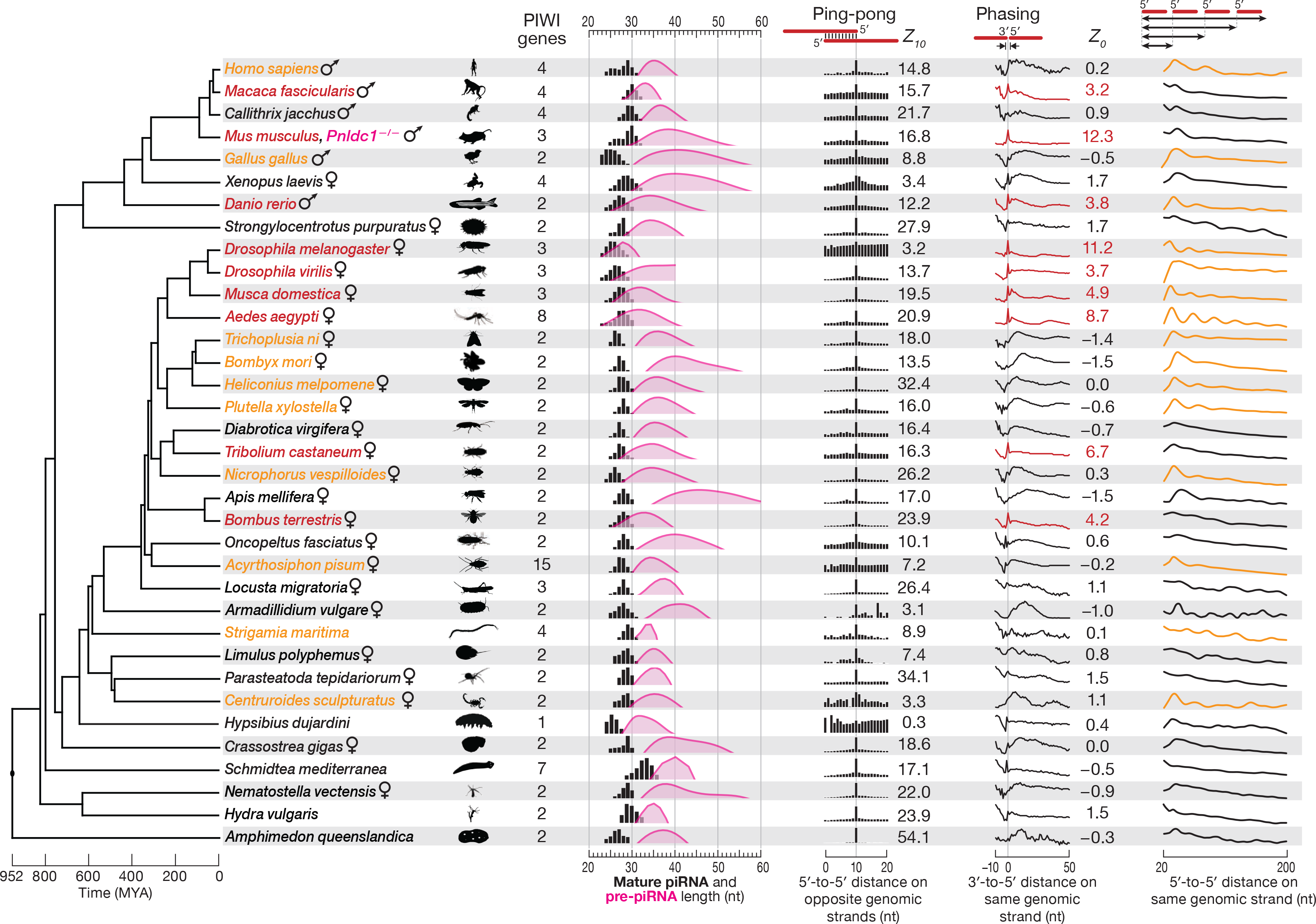
Evidence for Ping-Pong and Phased Pre-piRNAs among Animals. Cladogram of a representative set of 35 animal species showing the approximate time of each divergence. piRNA length profiles are shown for all sequencing reads taking into account their abundance; distance probability analyses are for ≥24-nt sequencing reads without taking into account abundance. Data for piRNA 5′-to-5′ distance was processed using non-parametric regression (LOWESS), and the first peak of the smoothed data was used to estimate pre-piRNA length. The significance of ping-pong (peak at 10 for 5′-to-5′ distance on opposite genomic strands) and phasing (peak at 0 for 3′-to-5′ distance on the same genomic strand) signatures was assessed using *Z-*scores. Red: species with *Z_0_* > 1.96 (*p* < 0.05) for 3′-to-5′ distance. Orange: species for which autocorrelation analysis detected periodic peaks of 5′-to-5′ piRNA distances. All data are from wild-type animals, except *Mus musculus* (*Pnldc1^−/−^*).

We estimated the length of pre-piRNAs in each of the 33 species by calculating the most frequent distance between mature piRNA 5′ ends (Figure 2). Consistently, if the estimated length range of pre-piRNAs overlapped the length range of mature piRNAs, we observed a tail-to-head arrangement of piRNAs: the most frequent distance between piRNA 3′ and 5′ ends was 0 (Figure 2). Strings of phased pre-piRNAs were detectable in an evolutionarily broad range of animals (red in Figure 2; *Z_0_* ≥ 1.96, i.e., *p* ≤ 0.05): the primate *Macaca fascicularis* (crab-eating macaque); the teleost fish *Danio rerio* (zebrafish); three dipteran insects, *Drosophila virilis*, *Musca domestica* (house fly), and *Aedes aegypti* (yellow fever mosquito); the coleopteran insect *Tribolium castaneum* (red flour beetle); and the hymenopteran insect *Bombus terrestris* (buff-tailed bumblebee). In other animals, pre-piRNA length was estimated to be significantly longer than that of mature piRNAs, suggesting that in these species, phased pre-piRNAs undergo extensive 3′-to-5′ trimming during maturation into primary piRNAs, making the detection of phasing infeasible in wild-type animals.

To test whether additional members of the group of 33 species used a phased primary biogenesis mechanism to produce piRNAs, we analyzed the 5′-to-5′ distances of mature piRNAs as a surrogate for detecting tail-to-head strings of pre-piRNAs (Figure S1). Such a strategy is expected to detect phasing when the distribution of pre-piRNA lengths is essentially unimodal. Species with phased piRNAs are predicted to display mature primary piRNAs whose 5′ ends are separated by multiples of the modal pre-piRNA length, yielding a repeating pattern of 5′-to-5′ piRNA distances (Figures 2 and S1). Autocorrelation analysis of the data allowed the quantitative detection of periodic peaks—evidence for phased pre-piRNA production—for ten additional animal species (orange in Figure 2; raw data in Figure S1): the arachnid *Centruroides sculpturatus* (bark scorpion); the centipede *Strigamia maritima*; the hemipteran insect *Acyrthosiphon pisum* (pea aphid); the coleopteran insect *Nicrophorus vespilloides* (burying beetle); four lepidopteran insects, *Trichoplusia ni* (cabbage looper), *Bombyx mori* (silk moth), *Heliconius melpomene* (postman butterfly), and *Plutella xylostella* (diamondback moth); the bird *Gallus gallus* (chicken); and the primate *Homo sapiens* (human).

Our analysis of silk moth adult ovary piRNAs as well as earlier reporter transgene experiments in cultured silk moth BmN4 cells (Homolka et al., 2015) suggest that the primary piRNA pathway collaborates with the secondary pathway to produce piRNAs in *B. mori*. However, a recent study reported that BmN4 piRNAs are made solely by the secondary pathway (Nishida et al., 2018). To further test the idea that *B. mori*, like the three other lepidopteran species we analyzed, produces phased primary piRNAs, we reanalyzed published sequencing data from BmN4 cells in which the piRNA maturation enzyme Trimmer/PNLDC1 was depleted by RNAi (Izumi et al., 2016). These data further support our conclusions: the most frequent 3′-to-5′ distance for piRNAs in Trimmer/PNLDC1 depleted BmN4 cells was zero (Figure S2A).

Finally, we detected significant piRNA ping-pong—the hallmark of the secondary piRNA biogenesis pathway—in 32 of the 33 non-model animals analyzed. Together, our data suggest that the secondary and primary piRNA biogenesis pathways likely collaborated to make piRNAs in the last common ancestor of all bilateral animals, an evolutionary distance of ~800 million years (Hedges et al., 2015; Kumar et al., 2017).

### *M. musculus* MILI and MIWI Participate in Phased Pre-piRNA Production

Our finding that phased primary piRNA production is a deeply conserved feature of piRNA biogenesis prompted us to reexamine how pachytene piRNAs are produced in the male germline of post-natal mice (*M. musculus*). The current model for piRNA biogenesis in post-natal male mouse germ cells presumes that mature piRNA length heterogeneity reflects different extents of PNLDC1-mediated trimming for MILI- and MIWI-bound pre-piRNAs (Figure 1, at left; Kawaoka et al., 2011; Izumi et al., 2016). The model predicts that (1) in the absence of the 3′-to-5′ trimming enzyme PNLDC1, piRNAs should be replaced by longer, untrimmed pre-piRNAs, and (2) untrimmed pre-piRNAs bound to MILI or MIWI should have similar length distributions (Figure 1, at left).

To test these predictions, we generated *Pnldc1^−/−^* mutant mice and sequenced small RNAs from post-natal spermatogonia, primary spermatocytes, secondary spermatocytes and round spermatids purified by fluorescent-activated cell sorting (FACS). Consistent with previous studies using whole testis (Ding et al., 2017; Zhang et al., 2017; Nishimura et al., 2018), the purified cell types all contained 25–40 nt small RNAs rather than the normal complement of mature 25–31 nt piRNAs (Figure 3A). We note that the length of *Pnldc1^−/−^* small RNAs is shorter than the pre-piRNA length estimated from the most frequent 5′-to-5′ distance of mature piRNAs (28–50 nt; Han et al., 2015). Nonetheless, our analyses show that the majority of *Pnldc1^−/−^* small RNAs are bona fide pre-piRNAs. First, the probability of 3′-to-5′ distances for *Pnldc1^−/−^* small RNAs peaked sharply at 0, the distance expected for pre-piRNAs which are produced tail-to-head, one after another (Figure 3A). Second, the genomic nucleotide immediately 3′ to *Pnldc1^−/−^* small RNAs was most often uridine (Figure 3A), consistent with both the 5′ U bias of primary piRNAs (Aravin et al., 2006; Girard et al., 2006) and with pre-piRNAs being produced tail-to-head.

**Figure 3.**
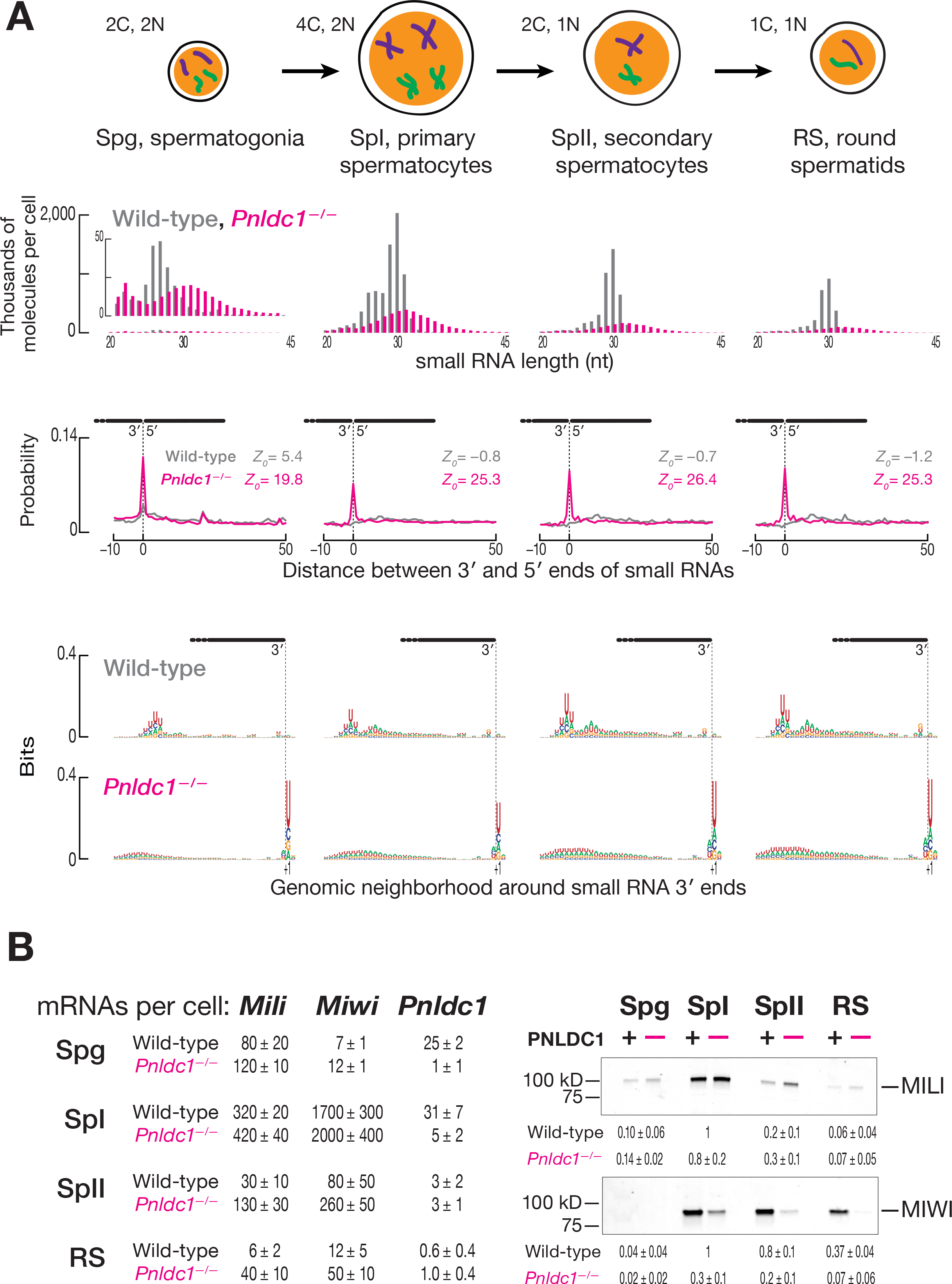
Mature piRNAs are Replaced by Untrimmed Pre-piRNAs in *Pnldc1^−/−^* Mice. (A) Male germ cell small RNA length profiles and, for ≥24-nt small RNAs, the probability of 3′-to-5′ distances on the same genomic strand and nucleotide composition of the genomic neighborhood of small RNA 3′ ends in wild-type and *Pnldc1^−/−^* mice. Data are from a single representative biological sample excluding reads with non-templated 3′ nucleotides. C, cell chromosome content; N, ploidy. (B) At left, mean (± standard deviation; *n* = 3) steady-state molecular abundance of *Mili*, *Miwi*, and *Pnldc1* mRNAs in male germ cells purified from wild-type and *Pnldc1^−/−^* mice. Spg, spermatogonia; SpI, primary spermatocytes; SpII, secondary spermatocytes; RS, round spermatids. At right, Representative Western blot images and relative mean (± standard deviation; *n* = 3) steady-state abundance of MILI and MIWI proteins. Each lane contained lysate from ~11,000 cells. Figures S2C and S2D show uncropped Western blot images.

Third, a pre-piRNA is expected to be followed tail-to-head by a long RNA not yet converted to a pre-piRNA, a pre-pre-piRNA. Such pre-pre-piRNAs would have 5′ monophosphorylated ends and be longer than pre-piRNAs. To identify pre-pre-piRNAs, we sequenced 5′ monophosphorylated, single-stranded RNA fragments ≥150 nt long from wild-type primary spermatocytes. As predicted for authentic pre-pre-piRNAs, the most likely distance from the 3′ ends of *Pnldc1^−/−^* small RNAs to the 5′ ends of the long, 5′ monophosphorylated RNAs was 0 (Figure S2B). We conclude that while most *Pnldc1^−/−^* small RNAs are bona fide pre-piRNAs, some likely correspond to pre-piRNAs trimmed by an exonucleolytic activity present in *Pnldc1^−/−^* mutant germ cells. To exclude these off-pathway pre-piRNA products, we confined our analyses here only to those *Pnldc1^−/−^* small RNAs that are followed tail-to-head by another small RNA. Such bona fide pre-piRNAs account for ~60% of all *Pnldc1^−/−^* small RNAs in primary spermatocytes.

The standard model for mouse post-natal piRNA biogenesis postulates that untrimmed pre-piRNAs bound to MILI or MIWI will have similar length distributions (Figure 1, at left). To test this prediction, we examined pre-piRNAs bound to MILI and MIWI in *Pnldc1^−/−^* primary spermatocytes—a cell type expressing both PIWI proteins (Figures 3B, S2C, and S2D; Table S1). Surprisingly, pre-piRNAs bound to MILI (mode = 31 nt) and MIWI (mode = 34 nt) had different length distributions (Figure 4A). How could a common machinery generate distinct lengths of pre-piRNAs for MILI and MIWI? One idea is that pre-piRNAs are sorted by length between the two PIWI proteins. A sorting model predicts that both short and long pre-piRNAs should be produced throughout spermatogenesis, regardless of presence of MILI or MIWI. Our data do not support a sorting model: the length of pre-piRNAs estimated from the most frequent 5′-to-5′ distance of wild-type piRNAs increases from 35 nt in spermatogonia, where only MILI is present, to 38 nt in primary spermatocytes, where both MIWI and MILI are expressed (Figure S2E).

**Figure 4.**
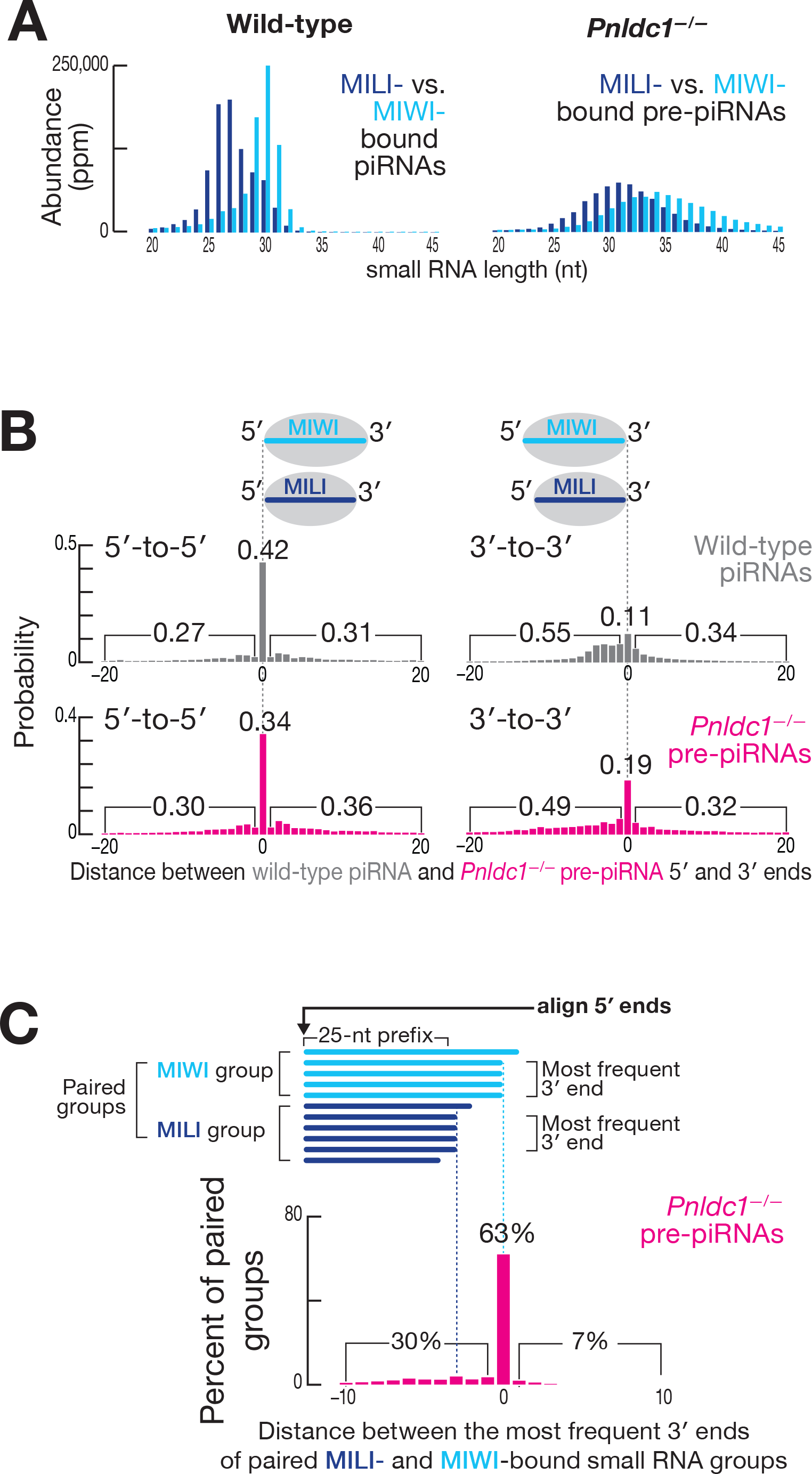
MILI and MIWI Participate in Phased Pre-piRNA Production. (A) Length profiles of MILI- and MIWI-bound piRNAs in wild-type and pre-piRNAs in *Pnldc1^−/−^* primary spermatocytes. Abundance was normalized to all genome mapping reads and reported in parts per million (ppm). Data are from a single representative biological sample; reads with non-templated 3′ nucleotides were excluded. (B) Probability of distances between the 5′ ends of MILI- and MIWI-bound piRNAs. Numbers indicate the total frequency of 5′ or 3′ ends of MILI-bound small RNAs residing before, after, or coinciding with the 5′ or 3′ ends of the MIWI-bound small RNAs. Data are from a single representative biological sample. (C) Distance between the most frequent 3′ end of the MILI-bound pre-RNA group and the most frequent 3′ end of the corresponding paired MIWI-bound pre-RNA group. Data are from a single representative biological sample.

Another explanation for the difference in the modal length of MILI- and MIWI-bound pre-piRNAs is that PIWI proteins themselves position the endonuclease that converts pre-pre-piRNAs to pre-piRNAs. This model predicts that a MILI-bound pre-piRNA should be shorter than a MIWI-bound pre-piRNAs when the two pre-piRNAs have the same 5′ end. To test this, we asked whether the genomic positions of the 5′ or 3′ ends of the pre-piRNAs bound to MILI differ from those of the pre-piRNAs bound to MIWI. For both piRNAs and pre-piRNAs, we calculated the probabilities of the 5′ ends of MILI-bound RNAs residing before, after, or coinciding with the 5′ ends of MIWI-bound RNAs. Similarly, we calculated the probabilities for MILI- or MIWI-bound piRNA and prepiRNA 3′ ends. In wild-type primary spermatocytes, MILI- and MIWI-bound mature piRNAs were more likely to share 5′ ends than 3′ ends (0.42 for 5′ ends vs. 0.11 for 3′ ends; Figure 4B). Similarly, in *Pnldc1^−/−^*, MILI- and MIWI-bound pre-piRNAs were more likely to share 5′ ends than 3′ ends (0.34 for 5′ ends vs. 0.19 for 3′ ends; Figure 4B). Consistent with the idea that the length of a mature piRNA is defined by the footprint of its PIWI partner (Kawaoka et al., 2011; Izumi et al., 2016), the 3′ ends of MILI-bound piRNAs were more likely to be upstream of the 3′ ends of MIWI-bound piRNAs in wild-type primary spermatocytes (0.55 upstream vs. 0.34 downstream; Figure 4B). Surprisingly, in *Pnldc1^−/−^* mutants, the probability of the 3′ ends of MILI-bound pre-piRNAs lying upstream of the 3′ ends MIWI-bound piRNAs was also higher (0.49 upstream vs. 0.32 downstream; Figure 4B). We obtained similar results analyzing only pre-piRNAs from pachytene piRNA loci—i.e., excluding those pre-piRNAs bound to MILI before the onset of meiosis (data not shown). Thus, MILI-bound pre-piRNAs are likely to share 5′ ends with MIWI-bound pre-piRNAs, but those pre-piRNAs bound to MILI are generally shorter than MIWI-bound pre-piRNAs.

These analyses compared all combinations of 3′ ends of pre-piRNAs, including those that do not share a common 5′ end. To examine the 3′ ends of pre-piRNAs produced from a common pre-pre-piRNA, we grouped together all unambiguously mapping pre-piRNAs with the same 5′, 25-nt prefix and used their read abundance to identify the most frequent 3′ end for each group (Figure S2F). We then paired the groups of MILI- and MIWI-bound wild-type piRNAs or *Pnldc1^−/−^* mutant pre-piRNAs that had the same 5′, 25-nt prefix. Analysis of the paired groups of MILI- and MIWI-bound pre-piRNAs showed that the most frequent 3′ end of the MILI-bound pre-piRNA group was either the same (63% of paired groups; Figure 4C) or upstream of the most frequent 3′ end of the corresponding MIWI-bound pre-piRNA group (30% of paired groups; Figure 4C). Thus, when MILI binds the 5′ end of a pre-pre-piRNA, it is rare for the resulting pre-piRNA to be longer than the pre-piRNA generated when MIWI binds the same pre-pre-piRNA (7% of paired groups; Figure 4C). Confining the analysis to pachytene piRNA loci produced identical results (data not shown).

We took advantage of the 5′ U bias of pre-piRNAs (Figure S3A) and the fact that pre-piRNAs are produced tail-to-head to further test the hypothesis that MILI and MIWI position the endonucleolytic cleavage that generates the 3′ ends of pre-piRNAs. Inspection of individual pre-piRNAs from pachytene piRNA loci suggested that when MILI- and MIWI-bound pre-piRNAs share the same 5′ end but differ in their 3′ ends, both 3′ ends are followed by a uridine in the genomic sequence (Figure 5, at left). For MILI- and MIWI-bound pre-piRNAs with identical 5′ and 3′ ends, the U following their shared 3′ end is the only uridine present in that genomic neighborhood.

To quantify these observations, we sorted the paired groups of pre-piRNAs bound to MILI and MIWI into cohorts according to the number of nucleotides separating the 3′ ends of the most frequent MILI- and MIWI-bound pre-piRNAs (Figure S2F). Thus, cohort 0 contained paired groups of pre-piRNAs whose most frequent 3′ end was identical for MILI- and MIWI-bound pre-piRNAs. In cohort 1, the most frequent MILI-bound pre-piRNA 3′ end lay 1 nt upstream of the 3′ end of the pre-piRNA bound to MIWI (Figure S2F). For each cohort, we measured the uridine frequency at each position in the genomic neighborhood of the 3′ ends. Strikingly, whenever two separate peaks of high uridine frequency resided in a genomic neighborhood, the most frequent 3′ end of MILI-bound pre-piRNA groups lay immediately before the upstream uridine, while the most frequent 3′ end of MIWI-bound pre-piRNA groups lay before the downstream uridine (Figure 5, at right). When a single peak of uridine was surrounded by a uridine desert, the 3′ ends of the MILI- and MIWI-bound pre-piRNAs coincided at the nucleotide immediately before the single uridine peak (Figure 5). We obtained essentially indistinguishable results analyzing all genome-mapping pre-piRNAs (data not shown).

**Figure 5.**
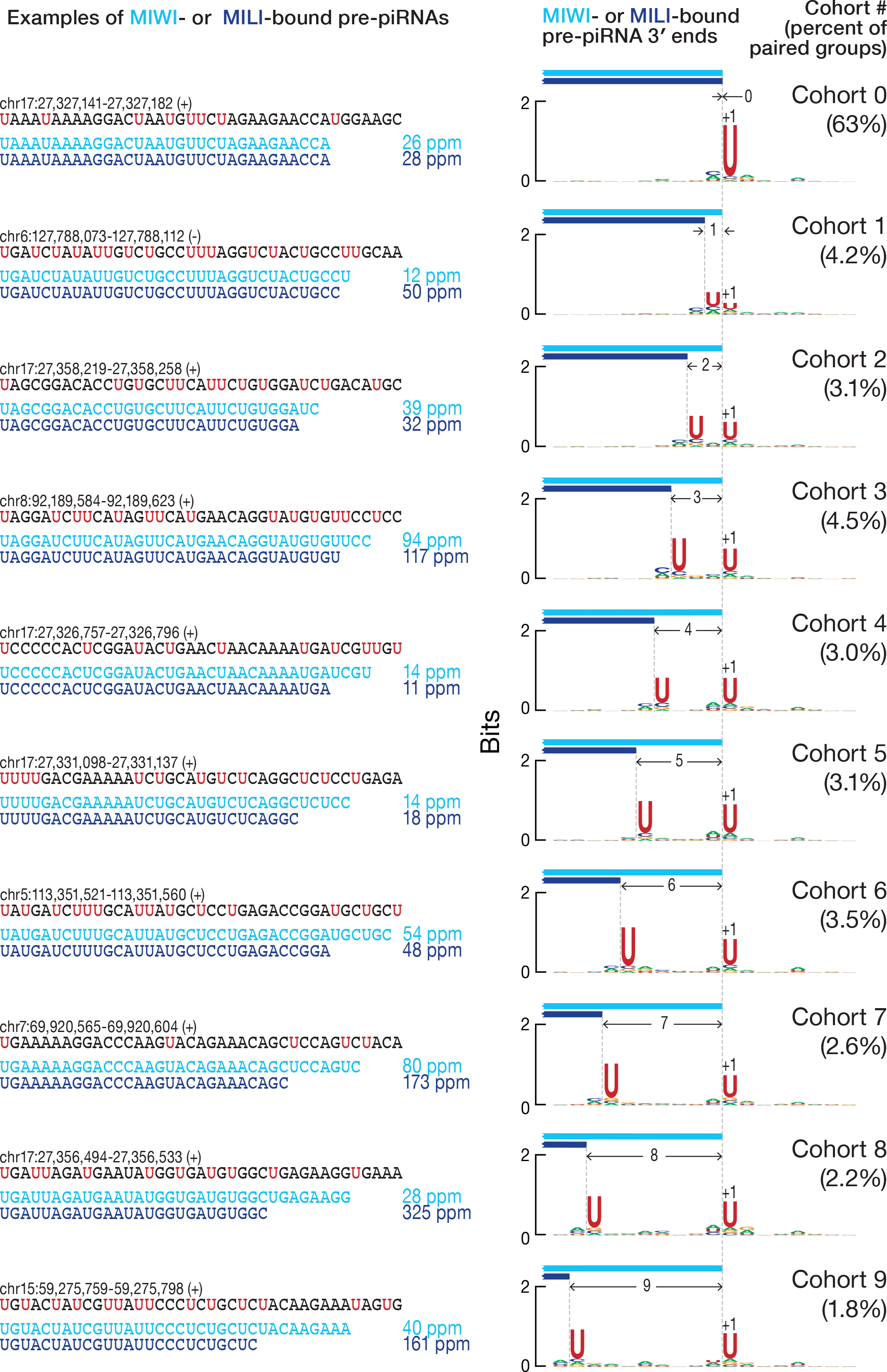
PIWI Protein Identity and the Availability of Uridines Dictate the Position of Mouse Pre-piRNA 3′ Ends. At left, a comparison of the most frequent 3′ ends of individual MILI- and MIWI-bound pre-piRNAs sharing the same 5′ end in *Pnldc1^−/−^* primary spermatocytes. Data are in parts per million (ppm) for pre-piRNAs derived from pachytene piRNA loci. At right, the nucleotide bias of the genomic neighborhood around the most frequent 3′ ends of paired MILI- and MIWI-bound pre-piRNA groups in *Pnldc1^−/−^* primary spermatocytes. Data are from a single representative biological sample.

Together, these data suggest that MILI and MIWI directly participate in the production of pre-piRNAs by positioning the endonuclease generating pre-piRNA 3′ ends.

### MILI and MIWI Bind a Pre-pre-piRNA 5′ End, then Position the Endonuclease

For the pachytene piRNA loci, 63% of paired groups of MILI- and MIWI-bound pre-piRNAs belonged to cohort 0 (Figure 4C). Even though a range of 3′ ends was present for these paired groups, the most frequent 3′ end for a MILI-bound pre-piRNA group was the same as the most frequent 3′ end for the corresponding MIWI-bound pre-piRNA group. In theory, this observation could reflect an underlying non-random distribution of uridines in piRNA precursor transcripts. Pachytene piRNA precursor transcripts are 27.4% U, which is expected for a near random uridine distribution. However, these transcripts could hypothetically contain many uridine-free regions, forcing the pre-pre-piRNA cleaving endonuclease to cut 5′ to the only uridine available in the genomic region. To test this idea, we used random sampling to estimate the probability of uridine surrounded by non-uridine stretches (VVVVUVVVV) and the probabilities of the two closest uridines separated by a given length of non-uridine sequence (UU, UVU, UVVU, UVVVU, etc.; Figure S3B). The data obtained from random sampling fit well to the geometric distribution, *P*(*x*) = *p*(100− *p*)^*x*−1^, where *p* = 27.4 (Pearson’s = 0.996; data not shown). Thus, uridines are spread randomly across the pachytene piRNA transcripts. Uridines were also found to be randomly distributed when the analysis was confined to the genomic neighborhoods around the paired groups of MILI- and MIWI-bound pre-piRNAs (data not shown). Therefore, the distribution of uridine in pachytene piRNA precursor transcripts cannot explain why, for 63% of paired groups of MILI- and MIWI-bound pre-piRNAs, their most frequent 3′ ends coincide (cohort 0 in Figure S3B). We conclude that the underlying mechanism of pre-piRNA production, not the sequence of pachytene piRNA precursor transcripts, determines the distribution of pre-piRNA 3′ ends.

If PIWI proteins position the endonuclease during piRNA precursor transcript fragmentation, they may do so by binding the 5′ end of pre-pre-piRNA. In this case, much of the sequence of the prospective pre-piRNA would be occluded by the PIWI protein and inaccessible for cleavage. Only uridines located 3′ to the PIWI protein’s footprint could be recognized by the endonuclease. Based on pre-piRNA length data, the footprint of MILI is expected to be ~3 nt smaller than that of MIWI (Figures S2E and 4A), giving the endonuclease access to more upstream uridines when it is positioned by MILI rather than MIWI. In this view, the endonuclease is constrained by the PIWI protein to cleave at the nearest uridine not masked by the protein’s footprint. Thus, when only a single exposed uridine is present locally, the resulting MILI-bound and MIWI-bound pre-piRNAs share a common 3′ end (cohort 0 in Figure 5).

This model makes two predictions. Imagine a MIWI-bound pre-piRNA corresponding to the shortest length permitted by MIWI’s footprint (Figure S3C). By definition, when this pre-piRNA is a member of cohort 0, the corresponding MILI-bound pre-piRNA has the same 3′ end—i.e., the two pre-piRNAs are identical. The model predicts that no uridines will be found in the last three nucleotides of the pre-piRNA, but that there may be uridines ≥4 nt upstream of the pre-piRNA 3′ end. These uridines are inaccessible to the endonuclease, because they are concealed within the footprint of MILI. In other words, because of MILI’s footprint, cohort 0 is expected to incorporate all instances in which uridines are ≥4 nt upstream of the uridine used by the endonuclease positioned by MIWI (Figure S3C). To test this prediction, we used uridines randomly sampled in pachytene piRNA clusters as the 5′ ends of simulated pre-piRNAs. We set the 3′ ends of the simulated pre-piRNAs immediately before the first uridine occurring >31 nt (simulated MILI footprint) or >34 nt (simulated MIWI footprint) downstream of the simulated pre-piRNA 5′ end (Figure S3D). We then sorted the pairs of simulated MILI- and MIWI-bound pre-piRNAs sharing their 5′ ends into cohorts based on the number of nucleotides separating their 3′ ends. The result of the simulation fit well to the biological data (Pearson’s ? = 0.94; Spearman’s ? = 0.92; Figure S3D) further supporting the idea that MILI and MIWI differentially direct endonucleolytic cleavage after they bind the prepre-piRNA 5′ end.

The model makes a second prediction: the MILI-bound pre-piRNAs present in cohorts 4 and greater (Figure 5) will be paired with atypically long MIWI-bound pre-piRNAs (Figure S3E). In fact, the median length of MIWI-bound pre-piRNAs in cohorts 4–9 was longer than that in cohorts 0–3 (35–38 nt vs. 32–34 nt; Figure S3F). Thus, the data support the idea that the footprints of MILI and MIWI restrict which uridines can be used to generate pre-piRNA 3′ ends.

We conclude that MILI and MIWI position the pre-pre-piRNA cleaving endonuclease when it establishes pre-piRNA 3′ ends. When PIWI proteins bind the 5′ ends of pre-pre-piRNAs to initiate this process, the distinct sizes of the MILI and MIWI footprints differentially restrict the uridines available to the endonuclease. Both MILI and MIWI direct the endonuclease to cleave 5′ to the nearest uridine, but MILI positions the endonuclease upstream of the site dictated by MIWI.

### The Number of piRNAs per Cell Corresponds to the Number of PIWI proteins

The model proposed here also posits that a PIWI protein must bind the 5′ end of a piRNA precursor before the pre-piRNA can be liberated and trimmed into a mature piRNA. Thus, the number of PIWI proteins per cell is expected to be similar to the number of mature piRNAs.

To quantify the abundance of piRNAs in FACS-purified mouse male germ cells, we added known amounts of 18 synthetic RNA oligonucleotides to the total mouse RNA before preparing sequencing libraries. By comparison with the synthetic oligoribonucleotides, we estimated that the number of 24–33-nt RNAs (i.e., mature piRNAs) in primary spermatocytes (7.8 ± 0.6 × 10^6^ RNAs/cell), secondary spermatocytes (3.9 ± 0.1 × 10^6^ RNAs/cell), and round spermatids (2.5 ± 0.2 × 10^6^ RNAs/cell) corresponds to 5.7 ± 0.2–7.2 ± 0.6 µM (Figure S4A). Notably, the 100 most abundant piRNA species are present at ~2,400–19,000 molecules per cell, comparable to the most abundant miRNAs in these cells: ~2,500 molecules of let-7a; ~2,500 molecules of miR-449a; and ~23,500 molecules of miR-34c per cell.

Next, we assessed the number of MILI and MIWI proteins in the FACS-purified germ cells by western blotting, using recombinant, SNAP-tagged PIWI proteins as concentration standards (Figures S4B and S4C). The total number of PIWI proteins per cell (5.5 ± 1.5 × 10^6^/cell in primary spermatocytes, 3.0 ± 0.4 × 10^6^/cell in secondary spermatocytes, 1.5 ± 0.5 × 10^6^/cell in round spermatids) correlated well with the total number of piRNAs at different stages (Pearson’s ? = 0.99), and the total cellular concentration of PIWI proteins (4 ± 1–4.8 ± 1 µM; Figure S4A) was similar to that of piRNAs (5.7 ± 0.2–7.2 ± 0.6 µM; Figure S4A).

If virtually all piRNAs are bound to a PIWI protein, the length distribution of total piRNAs in cells should be explained by the combination of MILI- and MIWI-bound piRNA lengths. In primary spermatocytes, a cell type expressing both PIWI proteins, the simulated length profile of total piRNAs fit well to the biological data when the ratio of MILI to MIWI was based on our estimate of their absolute abundance (16 MILI molecules per 84 MIWI molecules; Pearson’s = 0.99; Figure S4D).

Taken together, these data support a model in which piRNA biogenesis requires a PIWI protein to bind the 5′ end of each piRNA precursor transcript to initiate pre-piRNA production.

### *D. melanogaster* Piwi and Aub Also Participate in Phased Pre-piRNA Production

To test if PIWI-protein binding similarly positions the endonuclease that establishes the 3′ ends of pre-piRNAs in *D. melanogaster*, we analyzed data from fly ovaries. *D. melanogaster* makes three PIWI proteins—Piwi, Aubergine (Aub), and Argonaute3 (Ago3). Ago3 and Aub collaborate to generate secondary piRNAs via the ping-pong cycle, whereas Piwi and, to a lesser extent, Aub, bind primary piRNAs (Mohn et al., 2015; Han et al., 2015; Senti et al., 2015; Wang et al., 2015). The modal lengths of Piwi- and Aub-bound piRNAs differ by just one nucleotide (26 and 25 nt, respectively).

Fly piRNAs are less extensively trimmed than those in mice, so the sequences of mature piRNAs more readily reveal the mechanics of pre-piRNA biogenesis. Analysis of the 3′-to-5′ distance between mature piRNAs showed that Piwi- and Aub-bound piRNAs are often found tail-to-head in any combination of the two proteins (Figure S5A). In addition, the probability of common 5′ ends between Piwi- and Aub-bound piRNAs was higher than the probability of common 3′ ends (0.34 for 5′ vs. 0.20 for 3′ ends; Figure S5B). To analyze the 3′ ends of piRNAs produced from a common pre-pre-piRNA, we grouped Piwi- and Aub-bound piRNAs sharing a common 5′, 23-nt prefix. After pairing the Piwi and Aub groups, we found that the most frequent 3′ ends of Aub-bound piRNA groups often coincided (66% of paired groups; Figure S5C) or lay upstream (26% of paired groups; Figure S5C) of the most frequent 3′ ends of the corresponding Piwi-bound piRNA groups. For just 8% of paired groups, the most frequent 3′ ends of Aub-bound piRNAs were downstream of the most frequent 3′ ends of the corresponding Piwi-bound piRNAs.

Next, we divided the paired Piwi- and Aub-bound piRNA groups into cohorts according to the distance between the most abundant 3′ ends for Piwi- and Aub-bound piRNAs. When the most frequent 3′ ends of Piwi- and Aub-bound piRNA groups were identical, they lay immediately before a single uridine in a uridine-depleted genomic neighborhood (Figure 6A). Conversely, when there were two peaks of uridine, the most frequent 3′ ends of Aub-bound piRNA groups lay immediately before the upstream peak, while the most frequent 3′ ends of Piwi-bound piRNA groups were immediately before the downstream peak (Figure 6A). Confining the analyses to piRNAs arising from germline-specific transposons (Wang et al., 2015) produced essentially the same results (data not shown).

**Figure 6.**
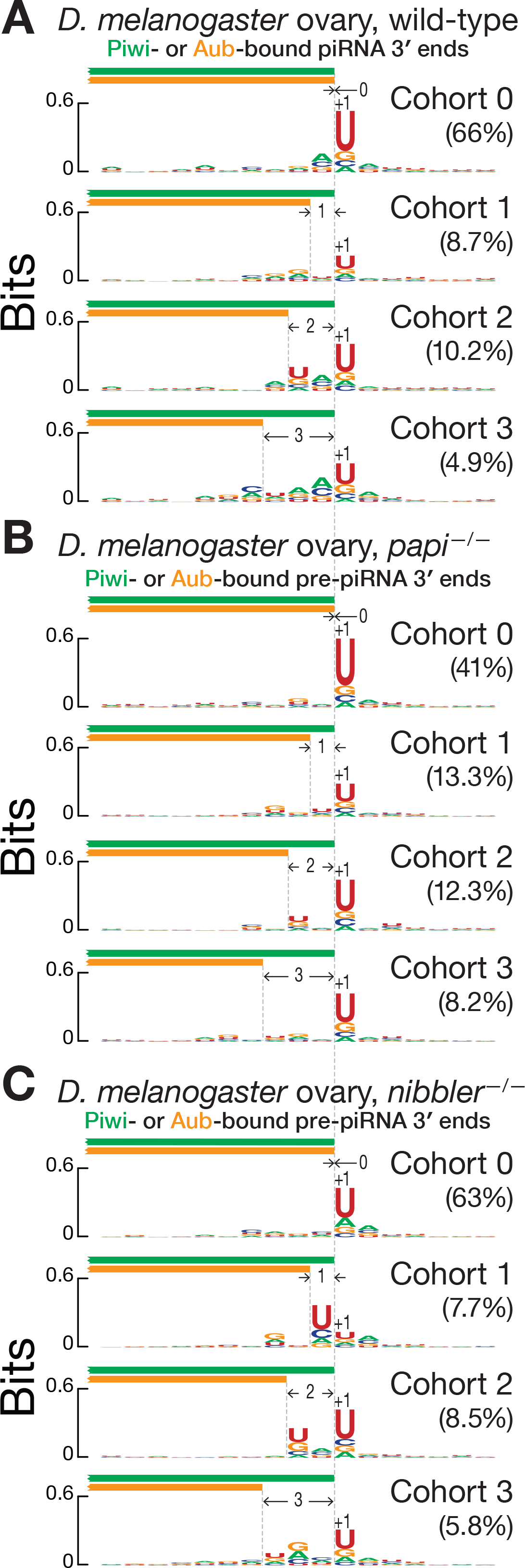
PIWI Protein Identity and the Availability of Uridines Dictate the Position of *Drosophila melanogaster* Pre-piRNA 3′ Ends. Nucleotide bias of the genomic neighborhood around the most frequent 3′ ends of paired Piwi- and Aub-bound piRNA groups from wild-type (A), and pre-piRNA groups in *papi^−/−^* (B) and *nibbler^−/−^* (C) *D. melanogaster* ovaries. Data are for all unambiguously mapping RNAs from a single representative biological sample.

Like mice, flies require Papi for pre-piRNA trimming: without Papi, fly pre-piRNAs have a median length 0.35 nt longer than wild-type piRNAs (Han et al., 2015; Hayashi et al., 2016). However, flies lack a PNLDC1 homolog (Hayashi et al., 2016). Instead, the miRNA-trimming enzyme Nibbler (Han et al., 2011; Liu et al., 2011) trims fly pre-piRNAs (Feltzin et al., 2015; Wang et al., 2016; Hayashi et al., 2016). In *papi^−/−^* and *nibbler^−/−^* mutant ovaries, Piwi- and Aub-bound pre-piRNAs are phased (Figure S5D). As in mice, phasing occurs between Aub- and Piwi-bound, Aub- and Aub-bound, Piwi- and Piwi-bound, and Piwi- and Aub-bound piRNAs, supporting the idea that a single pre-pre-piRNA molecule can generate both Aub- and Piwi-bound pre-piRNAs. Piwi- and Aub-bound pre-piRNAs are more likely to share 5′ ends than 3′ ends (*papi^−/−^*: 0.23 for 5′ ends vs. 0.09 for 3′ ends; *nibbler^−/−^*: 0.29 for 5′ ends vs. 0.15 for 3′ ends). Moreover, the 3′ ends of Aub-bound pre-piRNAs are more likely to be found upstream than downstream of the 3′ ends of Piwi-bound pre-piRNAs (*papi^−/−^*: 0.51 upstream vs. 0.40 downstream; *nibbler^−/−^*: 0.49 upstream vs. 0.36 downstream).

Grouping and pairing Piwi- and Aub-bound pre-piRNAs with common 5′, 23-nt prefix revealed that the most frequent 3′ end of Aub-bound pre-piRNA groups either coincided with the most frequent 3′ end of the corresponding Piwi-bound pre-piRNA group (*papi^−/−^*: 41% of paired groups; *nibbler^−/−^*: 63% of paired groups) or lay upstream of the most frequent 3′ end of the Piwi-bound pre-piRNA group (*papi^−/−^*: 43% of paired groups; *nibbler^−/−^*: 28% of paired groups). For only a small minority of paired groups are Aub-bound pre-piRNAs longer than Piwi-bound pre-piRNAs (*papi^−/−^*: 16% of paired groups; *nibbler^−/−^*: 9% of paired groups). As in mice, when a fly genomic neighborhood contained a single uridine frequency peak, the most frequent 3′ ends of Piwi- and Aub-bound pre-piRNA groups were identical and mapped immediately before the peak (Figures 6B and 6C): i.e., the most frequent Piwi- and Aub-bound pre-piRNAs had the same sequence and genomic coordinates. In contrast, when two uridine frequency peaks were present, the most frequent 3′ ends of Aub-bound pre-piRNA groups preceded the upstream peak, while the most frequent 3′ ends of the corresponding Piwi-bound pre-piRNA groups lay before the downstream peak.

Collectively, these data suggest that, as for MILI and MIWI in mice, Aub and Piwi in flies position the endonuclease that simultaneously generates the pre-piRNA 3′ end and the 5′ end of the succeeding pre-pre-piRNA. Thus, mammalian and insect PIWI proteins directly participate in establishing the pattern of phased pre-piRNA biogenesis.

### MILI and MIWI Slicing of Long piRNA Precursor Transcripts Initiates Pre-Pre-piRNA Biogenesis in Post-Natal Mice

Pre-piRNAs appear to be less stable than mature piRNAs in post-natal mice: primary spermatocytes, secondary spermatocytes and round spermatids from *Pnldc1^−/−^* testes contain approximately 2–3 times less 24–45 nt RNAs than wild-type cells, whereas miRNA abundance is unchanged (Figure S4A). Although piRNA cluster transcript abundance, measured by RNA-seq and RT-qPCR of RNA from whole testes, has been reported to be unchanged in *Pnldc1^−/−^* mice (Zhang et al., 2017), our RNA-seq data from FACS-purified germ cells show that without PNLDC1, the steady-state levels of many piRNA cluster transcripts increase. Absolute transcript abundance increased both in secondary spermatocytes (median increase = 3.0×; FDR ≤ 0.1) and round spermatids (median increase = 4.5×; FDR ≤ 0.1) for the ten loci that produce the most pachytene piRNAs and which account for ~50% of all piRNAs in meiotic and post-meiotic wild-type cells. At the same time, the absolute abundance of piRNAs from these loci decreased both in secondary spermatocytes (median decrease = 2.5×; FDR ≤ 0.1) and round spermatids (median decrease = 3.7×; FDR ≤ 0.1).

A possible explanation for the increased steady-state levels of pachytene piRNA cluster transcripts and the decreased amount of pre-piRNAs in *Pnldc1^−/−^* mutants is that piRNAs themselves are required to process piRNA cluster transcripts into phased pre-piRNAs. Indeed, in the *D. melanogaster* female germline, biogenesis of primary phased piRNAs is initiated by a slicing event directed by an Ago3-bound secondary piRNA generated via the ping-pong amplification pathway (Mohn et al., 2015; Han et al., 2015; Senti et al., 2015; Wang et al., 2015). Thus, secondary piRNAs initiate phased primary piRNA production by cleaving a piRNA cluster transcript to generate a pre-pre-piRNA. Although MILI initiates phased piRNA production in the neo-natal mouse testis (Yang et al., 2016), such a mechanism is not thought to play a role in piRNA biogenesis in the post-natal germline of male mice. We _reexamined this presumption._

To identify the pre-pre-piRNAs from which phased pre-piRNAs are generated in mouse primary spermatocytes, we selected from our data set of ≥150-nt, 5′ monophosphorylated RNA sequences from wild-type cells, those RNAs derived from the pachytene piRNA loci. Many of these shared 5′ ends with mature piRNAs, consistent with their corresponding to pre-pre-piRNAs (Figure S6A; 79% shared 5′ ends with MILI-bound mature piRNAs, 77% shared 5′ ends with MIWI-bound mature piRNAs). We separated the putative pre-pre-piRNAs—the long RNAs sharing their 5′ end with a mature piRNA—from those for which no piRNA with the same 5′ end could be found (control; Figure 7A). As expected for precursors of pre-piRNAs, most putative pre-pre-piRNAs began with uridine (65% of pre-pre-piRNAs sharing 5′ ends with MILI-bound piRNAs, 67% of those sharing 5′ ends with MIWI-bound piRNAs; Figure 7B). Unexpectedly, putative pre-pre-piRNAs also showed a significant enrichment for adenine at their tenth nucleotide (39% of pre-pre-piRNAs sharing 5′ ends with MIWI-bound piRNAs and 38% of those sharing 5′ ends with MILI-bound piRNAs; Figure 7B). Such a 10A bias is the hallmark of PIWI-protein catalyzed target slicing and reflects the intrinsic preference of some PIWI proteins for adenine at the t1 position of their target RNAs (Wang et al., 2014; Matsumoto et al., 2016).

**Figure 7.**
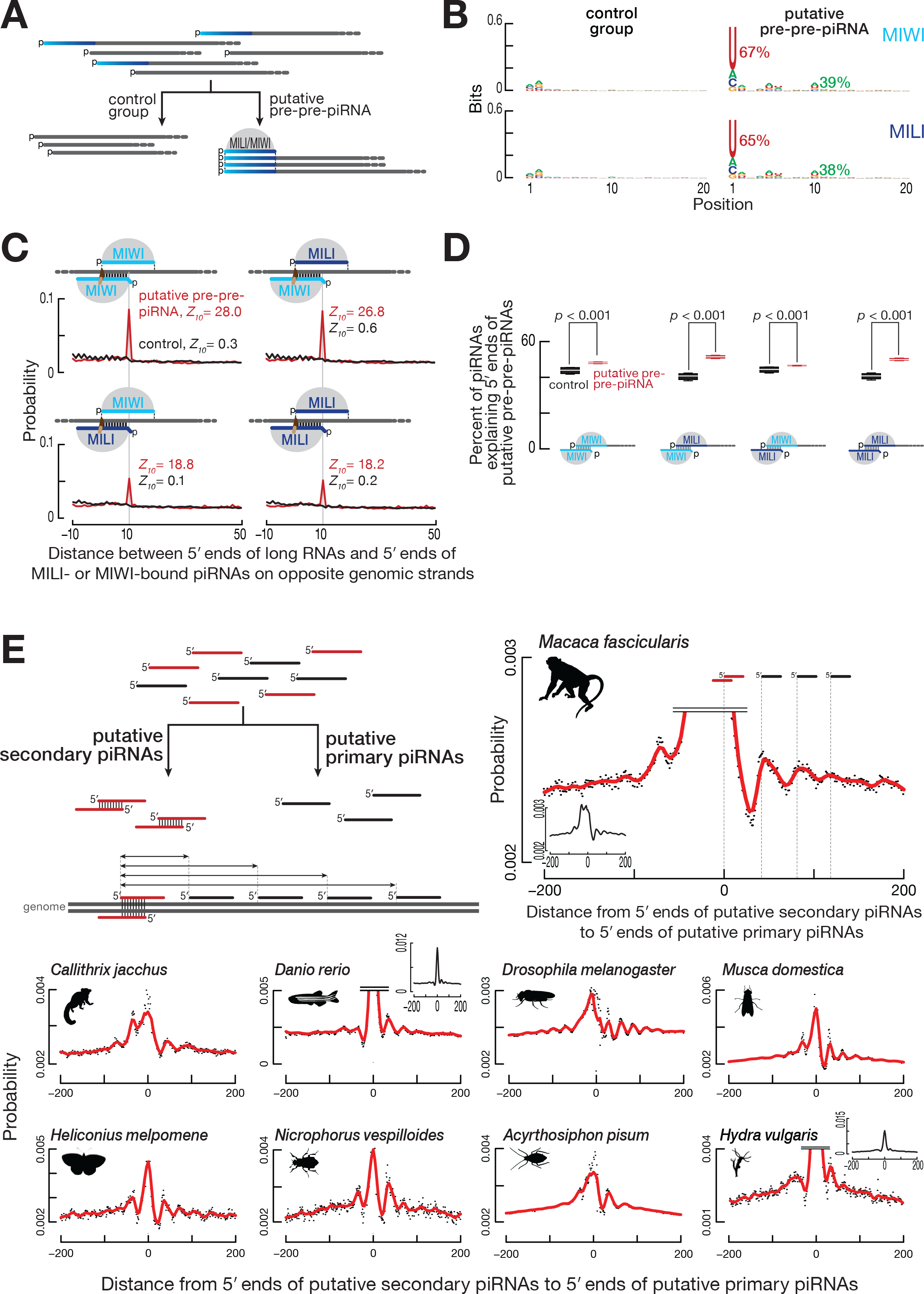
piRNA-directed, PIWI Protein-Catalyzed Slicing Initiates piRNA Biogenesis in Mice. (A) Strategy for the identification of putative pre-pre-piRNAs among ≥150-nt long, 5′ monophosphorylated RNAs derived from pachytene piRNA loci from wild-type mouse primary spermatocytes. RNAs sharing 5′ ends with MILI- or MIWI-bound mature piRNAs were classified as putative pre-pre-piRNAs; the remaining long RNAs served as the control group. Data in (B) and (C) are from a single representative biological sample. (B) Nucleotide bias of the 5′ ends of putative pre-pre-piRNAs compared to the control group. (C) Probability of distances between the 5′ ends of MILI- or MIWI-bound piRNAs and the 5′ ends of putative pre-pre-piRNAs or control RNAs complementary to nucleotides 2 to 10 of the piRNA (g2–g10). (D) Percent of piRNAs explaining the 5′ ends of either putative pre-pre-piRNAs or control RNAs. Nucleotides 2 to 10 of guide piRNAs (g2–g10) were required to be complementary to the target long RNAs. Data are for long RNAs and MILI- or MIWI-bound piRNAs derived from pachytene piRNA loci. All possible pairwise combinations of data sets from two biological replicates were used to calculate medians. Whiskers correspond to minimum and maximum values. Wilcoxon rank-sum test was used to assess statistical significance. (E) Strategy to identify putative secondary and primary piRNAs, and to measure the probability of distances from the 5′ ends of putative secondary piRNAs to the 5′ ends of putative primary piRNAs. Distance probability analyses are for ≥24-nt sequencing reads without taking into account abundance. Data was processed using non-parametric regression (LOWESS).

To test the idea that MILI- or MIWI-bound piRNAs direct slicing of piRNA cluster transcripts to generate pre-pre-piRNAs, we searched for MILI- and MIWI-bound piRNAs that could have guided production of the 5′ ends of the putative pre-pre-piRNAs. Unlike siRNAs and miRNAs, the extent of base-pairing required between a piRNA and its target RNA to support PIWI-protein catalyzed target slicing is incompletely understood. Minimally, nucleotides 2 to 10 of the guide piRNA (g2–g10) are expected to pair with the target RNA (Reuter et al., 2011; Gou et al., 2014; Goh et al., 2015; Zhang et al., 2015). We observed a statistically significant overlap between nucleotides g2–g10 of MILI- and MIWI-bound piRNAs and putative pre-pre-piRNAs, but detected no such overlap with the control RNAs (Figures 7C and 7D). This result is unlikely to reflect canonical *cis* ping-pong, in which the piRNA and target overlap in the genome sequence: ≫99% of piRNAs contributing to the overlap do not map to the same genomic location as their targets. That is, the majority of piRNAs act in *trans* through partial complementarity to cleave targets transcribed from a piRNA-producing locus different from their own.

To account for variance in sample sizes and the different contributions of species with high and low read abundance, we repeated the analyses using either data subsampling or random resampling of the data with replacement (bootstrapping). These analyses confirmed the presence of the ping-pong signature for the putative pre-pre-piRNAs but not for the control RNAs (Figures S6B and S6C).

### Transposon-Derived piRNAs Direct Pre-Pre-piRNA Biogenesis in Post-Natal Mice

Transposons are a possible source of piRNAs capable of targeting piRNA precursor transcripts, because the sequence of a transposon can potentially bear short but fruitful stretches of complementarity with other genomic copies of the same or a related transposon family in another piRNA locus. For the pachytene piRNA loci, we found that more than twice as many transposon-derived piRNAs contributed to the overlap with putative pre-pre-piRNAs than expected by chance: 34.2% observed vs. 16.9% expected, based on the fraction of transposon-mapping piRNAs for MILI-bound piRNAs, and 46.1% observed vs. 20.1% expected for MIWI-bound piRNAs. This finding is particularly striking given that the transcribed regions of the pachytene piRNA loci contain fewer transposon sequences than the genome as a whole (31.9% vs. 41.9%).

Collectively, these data show that PIWI slicing plays a central role in initiating phased primary piRNA biogenesis in animals as evolutionarily distant as flies and mice. They also suggest that transposon-derived piRNAs can function in the production of piRNAs not participating in transposon silencing.

### PIWI Slicing Initiates Phased Primary piRNA Biogenesis in Most Animals

Mice and flies are separated by ~800 million years of evolution, and yet both species use secondary piRNAs to direct PIWI slicing to initiate the production of primary piRNAs. We explored whether the same strategy is employed for piRNA production in other animals. Because there are no available data sets of 5′ monophosphorylated prepiRNAs or pre-pre-RNAs from non-model organisms, we used an alternative approach. We assumed that piRNAs with a ping-pong partner on the opposite genomic strand were produced by PIWI slicing (putative secondary piRNAs) and that the remaining piRNAs were generated by phased primary piRNA production (putative primary piRNAs; Figure 7E). For each of 33 non-model animals, we grouped piRNAs as either putative secondary or primary piRNAs. If PIWI slicing initiates phased piRNA production, the most frequent position of the 5′ ends of putative primary piRNAs is predicted to lie at regular intervals downstream of the 5′ ends of putative secondary piRNAs.

We detected periodic peaks of putative primary piRNA 5′ ends downstream of the 5′ ends of putative ping-pong pairs in data from 12 species (Figure 7E): the primates *Macaca fascicularis* (crab-eating macaque) and *Callithrix jacchus* (white-tufted-ear marmoset); the teleost fish *Danio rerio* (zebrafish); two dipteran insects *Musca domestica* (house fly) and *Aedes aegypti* (yellow fever mosquito); three lepidopteran insects, *Trichoplusia ni* (cabbage looper), *Heliconius melpomene* (postman butterfly), and *Plutella xylostella* (diamondback moth); the coleopteran insect *Nicrophorus vespilloides* (burying beetle); the hemipteran insect *Acyrthosiphon pisum* (pea aphid); the flatworm *Schmidtea mediterranea* (freshwater planarian); and the cnidarian *Hydra vulgaris* (fresh-water polyp). Together with flies and mice, we detect secondary piRNA-initiated phased primary piRNA production in 14 species spanning four phyla—Cnidaria, Platyhelminthes, Arthropoda, and Chordata—including four vertebrates and eight insects. Together, these data suggest that (1) the secondary pathway initiates phased primary piRNA production in species representing the major animal phyla including non-bilateral animals and (2) the last common ancestor of all animals produced piRNAs very much as animals make them today.

### Origins of the 5′ U Bias of Primary piRNAs

The 5′ U bias of primary piRNAs is thought to arise from the specificity of the pre-pre-piRNA cleaving endonuclease producing phased pre-piRNAs. However, the 5′ U bias could also reflect the preference of PIWI proteins to bind guide RNAs beginning with uridine (Kawaoka et al., 2011; Matsumoto et al., 2016). If PIWI proteins contribute to 5′ U bias of primary piRNAs, then selection for initial uridines should occur when MILI or MIWI binds the 5′ end of a pre-pre-piRNA. The frequency of 5′ U for pre-piRNAs bound to MILI or MIWI and sharing their 5′ ends with putative pre-pre-piRNAs is 91%, higher than the frequency of 5′ U for putative pre-pre-piRNAs (65% for those sharing 5′ ends with MILI-bound pre-piRNAs, 67% for those sharing 5′ ends with MIWI-bound pre-piRNAs). These data suggest that not all pre-pre-piRNAs are converted to pre-piRNAs with equal efficiency.

To determine whether first nucleotide identity influences the potential of pre-pre-piRNAs to produce pre-piRNAs, we asked if highly abundant pre-piRNAs were more likely to derive from pre-pre-piRNAs beginning with uridine. First, we sorted pre-piRNA species by their abundance into 10 equally sized bins. We then calculated the percent 5′ U for the putative pre-pre-piRNAs sharing their 5′ ends with the pre-piRNAs in each bin. For each bin, we also determined the ratio of pre-piRNA abundance to the abundance of the corresponding pre-pre-piRNAs. Consistent with the idea that the binding preference of PIWI proteins contributes to the 5′ uridine bias of piRNAs, putative pre-pre-piRNAs were more likely to produce pre-piRNAs when the pre-pre-piRNAs started with uridine (Figure S6D).

Notably, the frequency of 5′ U for mature piRNAs bound to MILI or MIWI and sharing their 5′ ends with putative pre-pre-piRNAs was the same as for untrimmed pre-piRNAs (91%). That no change in 5′ U bias is introduced at the trimming step agrees well with the current piRNA maturation model in which trimming of pre-piRNAs occurs after their loading into PIWI proteins.

## DISCUSSION

The analyses presented here suggest a unified model for piRNA biogenesis in the germ cells of mammals, insects and probably most other animals (Figure 1, at right). In the male germline of mice and the female germline of flies, PIWI proteins initiate piRNA biogenesis: PIWI-catalyzed, piRNA-guided slicing of a long piRNA precursor transcript creates a pre-pre-piRNA, whose monophosphorylated 5′ end serves as an entry point for further pre-piRNA production (Mohn et al., 2015; Han et al., 2015; Senti et al., 2015; Wang et al., 2015; Yang et al., 2016).

PIWI proteins also control downstream primary piRNA biogenesis: a PIWI protein bound to the pre-pre-piRNA 5′ end positions an endonuclease—perhaps Zucchini/PLD6—to carry out stepwise fragmentation of the pre-pre-piRNA into phased pre-piRNAs. The essential role of 5′ phosphate recognition for loading guide RNAs into PIWI and other Argonaute proteins (Nykanen et al., 2001; Ma et al., 2005; Parker et al., 2005; Wang et al., 2009; Frank et al., 2010; Boland et al., 2011; Kawaoka et al., 2011; Elkayam et al., 2012; Schirle and MacRae, 2012; Schirle et al., 2014; Cora et al., 2014; Wang et al., 2014; Matsumoto et al., 2016) makes it unlikely that a PIWI protein can bind a pre-pre-piRNA downstream of its 5′ end. Binding of a PIWI protein to the newly generated pre-pre-piRNA 5′ end defines the 3′ end of the prospective pre-piRNA: the endonuclease cleaves 5′ to the first accessible uridine following the footprint of the PIWI protein. This cut simultaneously creates the 3′ end of the pre-piRNA and 5′ end of the next pre-piRNA. Our data suggest that the 5′ U bias of phased pre-piRNAs reflects the combined action of the endonuclease cleaving 5′ to uridines and the preference of PIWI proteins for pre-pre-piRNAs beginning with uridine.

Collectively, the proposed model explains how (1) PIWI-protein slicing initiates and (2) PIWI-protein binding controls the production of phased pre-piRNAs. The resulting phased pre-piRNAs, still bound to PIWI proteins, are then trimmed to their characteristic length and 2′-*O*-methylated, generating a mature, functional piRNA ready to guide the PIWI protein. In some animals trimming plays a minor role in piRNA maturation, allowing their tail-to-head arrangement to be detected in the wild type, i.e., without removing the pre-piRNA trimming enzyme (Figure 2; Mohn et al., 2015; Han et al., 2015; Senti et al., 2015; Wang et al., 2015; Hayashi et al., 2016). Contrary to the conclusions of the recently published work by Nishida et al. (Nishida et al., 2018), our analyses revealed the presence of phased pre-piRNAs in an evolutionarily broad range of animals including Lepidoptera. Our data also argue against the idea that only the transcriptional silencing machinery relies on phased pre-piRNA production (Nishida et al., 2018). First, two cytoplasmic PIWI proteins bind phased pre-piRNAs in post-natal mouse testis. Second, among those animals we examined, phased pre-piRNAs were detected in 10 of the 20 animals possessing just two PIWI genes (Figure 2).

Although the revised model explains the data presented here, the details of individual steps remain to be established. Assigning specific functions to all the proteins known to act in processing single-stranded piRNA precursor transcripts into mature piRNAs remains a formidable challenge. Our model no doubt underestimates the complexity of piRNA biogenesis, whose individual steps occur in at least two cellular compartments: perinuclear nuage and mitochondria. We also do not understand what determinants destine pachytene piRNA cluster transcripts in mice, which are generated by conventional RNA Pol II transcription from euchromatic loci, to become piRNAs. One possible explanation is that enrichment of pachytene piRNA cluster transcripts with piRNA target sites could render them more likely targets of PIWI slicing.

The data presented here establish the essential role of PIWI-clade Argonautes in the generation of their own guide piRNAs and suggest a possible evolutionary trajectory for the piRNA pathway (Figure S7): PIWI-clade Argonautes emerged first, generating their own guides via the ping-pong cycle, perhaps assisted by co-opting a 3′-to-5′ exonuclease to trim the guide piRNA to a length more fully protected by the PIWI protein (Nishida et al., 2018). Later in evolution, PIWI proteins came to be aided by an endonuclease specialized for producing phased pre-piRNAs, allowing the production of additional piRNAs from the otherwise discarded 3′ product generated by the production of secondary piRNAs.

The Nematoda represent a particularly bizarre branch in the evolution of the piRNA pathway, because most lineages of this phylum appear to have lost PIWI proteins and piRNAs altogether (Wang et al., 2011; Sarkies et al., 2015; Holz and Streit, 2017). Remarkably, the PIWI-protein-producing nematode *Caenorhabditis elegans*, generates each 21U-RNA, as worm piRNAs are called, from a separate transcript. It is not known what mechanism generates the 5′ ends of 21U-RNAs, although the downstream steps of piRNA biogenesis are likely similar to animals of other phyla: loading of 21U-RNA precursor into PRG-1, 3′-to-5′ trimming to establish mature piRNA length (Tang et al., 2016), and 2′-*O*-methylation of the piRNA 3′ end (Kamminga et al., 2012; Montgomery et al., 2012; Billi et al., 2012).

The most diverged components of the piRNA pathway act in piRNA precursor transcription, while downstream proteins are conserved (Grimson et al., 2008; Klattenhoff et al., 2009; Handler et al., 2011; Cecere et al., 2012; Li et al., 2013; Hayashi et al., 2016; Andersen et al., 2017). Such deep conservation suggests that the core of the piRNA-producing machinery—with the secondary pathway (ping-pong) initiating the production of phased primary pre-piRNAs—probably appeared prior to the divergence of most animal lineages. Thus, despite differences among animals in the sources and functions of piRNAs, piRNAs are produced by a common strategy that reflects a common descent of the underlying machinery in metazoan evolution.

## ACCESSION NUMBERS

Sequencing data are available from the National Center for Biotechnology Information Small Read Archive using accession number PRJNA421205.

## AUTHOR CONTRIBUTIONS

I.G., C.C., A.A., and P.D.Z. conceived and designed the experiments. I.G., C.C., A.A., and K.C. performed the experiments. I.G. analyzed the sequencing data. I.G., C.C., and P.D.Z. wrote the manuscript.

## ACKNOWLEDGEMENTS

We thank S. Pechhold, T. Giehl, B. Gosselin, Y. Gu, and T. Krumpoch at UMass FACS Core for help sorting mouse germ cells; J. Gosselin and J. Gallant at UMass Transgenic Animal Modeling Core for help generating *Pnldc1^−/−^* mice; A. Boucher, C. Tipping, and G. Farley for technical assistance; Z. Weng, Y. Fu, and members of the Zamore laboratory for discussions and critical comments on the manuscript. This work was supported in part by National Institutes of Health grants GM65236 and P01HD078253 to P.D.Z.

## SUPPLEMENTAL FIGURE TITLES AND LEGENDS

**Figure S1, related to Figure 2.**
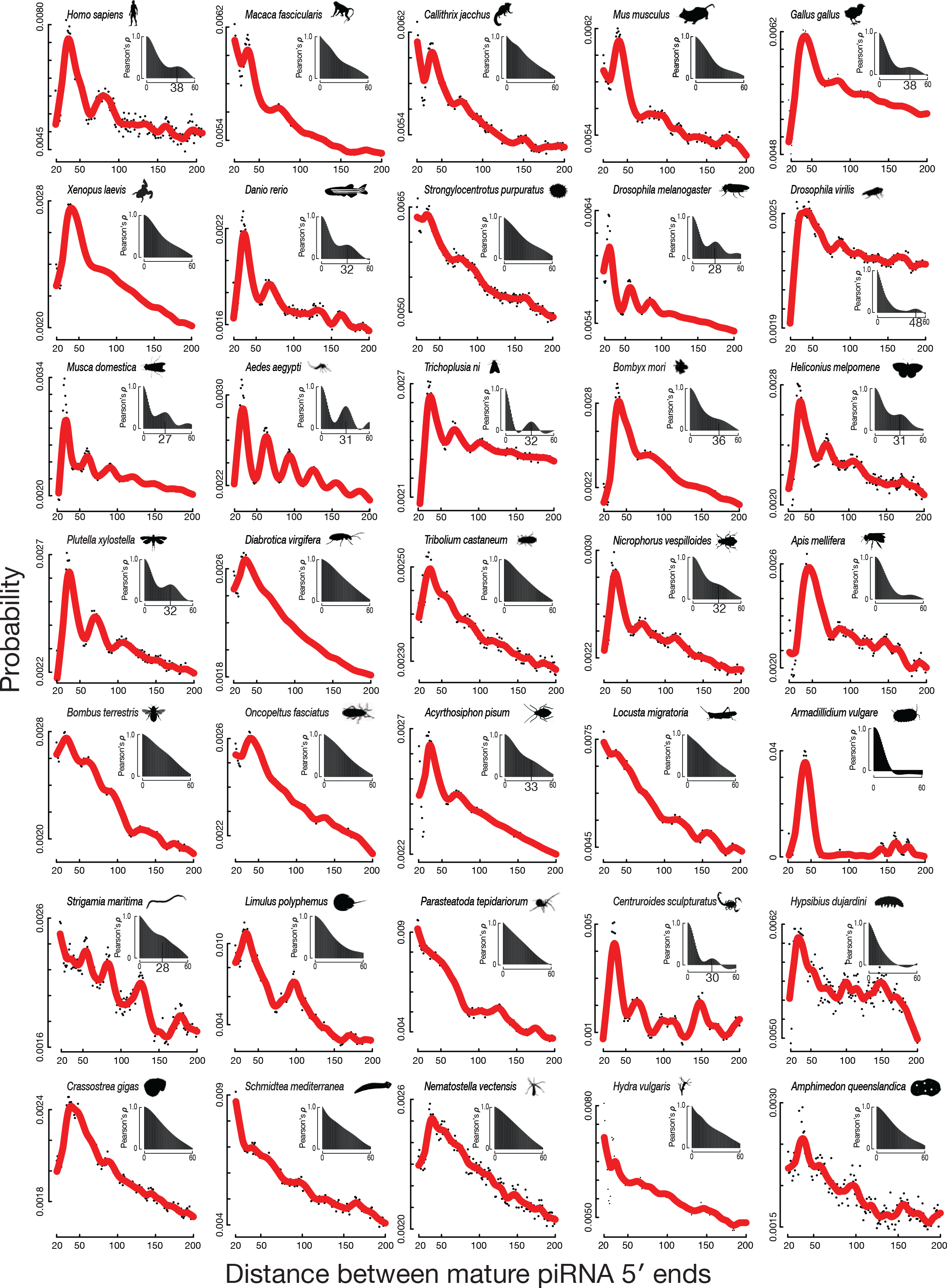
Probability of piRNA 5′-to-5′ distances on the same genomic strand for a representative set of 35 animal species. Distance probability analysis was done for ≥24-nt sequencing reads without taking into account abundance. Red: data smoothed using non-parametric regression (LOWESS). The insets show autocorrelation analysis of the smoothed data. The periodicity interval (i.e., estimated pre-piRNA length) was identified either as the peak of the autocorrelation function or the length closest to the zero data point of the derivative of the autocorrelation function. All data are from wild-type animals, except for *Mus musculus* (*Pnldc1^−/−^*).

**Figure S2, related to Figures 3 and 4.**
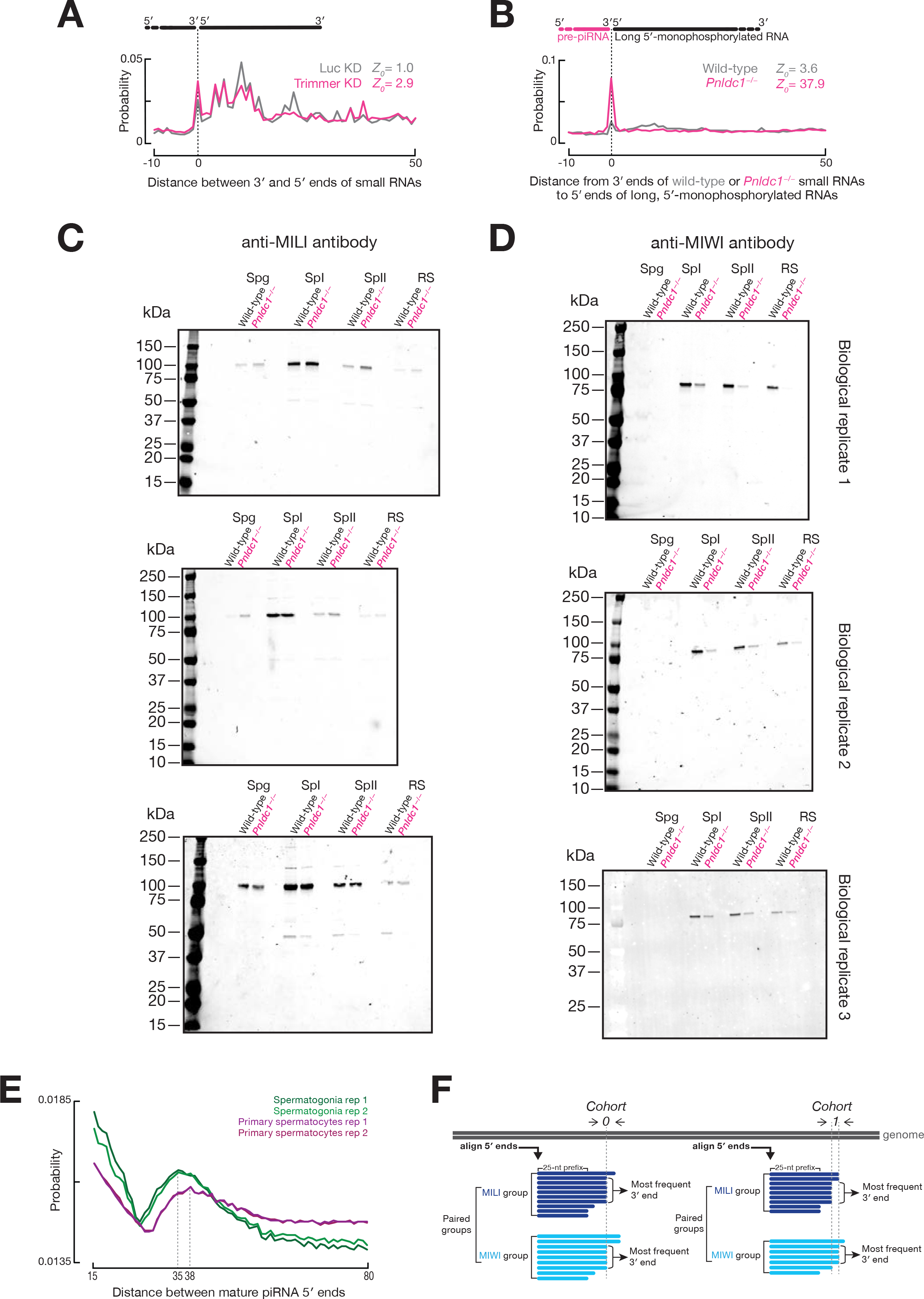
(A) Probability of 3′-to-5′ distances on the same genomic strand for ≥24-nt small RNAs in BmN4 cells. Either Trimmer/PNLDC1 (piRNA trimming enzyme) or *Renilla* luciferase (control) were depleted using RNAi (Izumi et al., 2016). (B) Probability of distances from the 3′ ends of ≥24-nt small RNAs in primary spermatocytes of wild-type or *Pnldc1^−/−^* mice to the 5′ ends of ≥150 nt long 5′ monophosphorylated RNAs in primary spermatocytes of wild-type mice. Data are from a single representative biological sample. (C, D) Relative abundance of MILI (C) and MIWI (D) in male germ cells wild-type and *Pnldc1^−/−^* mice assessed by Western blotting. Spg, spermatogonia; SpI, primary spermatocytes; SpII, secondary spermatocytes; RS, round spermatids. Each lane contains lysate from ~11,000 cells. Data are from three biological samples. (E) Probability of 5′-to-5′ distances on the same genomic strand for ≥24-nt small RNAs in spermatogonia and primary spermatocytes of wild-type mice. Data are from two biological samples. (F) Strategy to analyze 3′ ends of pre-piRNAs deriving from the same pre-pre-piRNA. First, pre-piRNAs with common 5′, 25-nt prefix are grouped and the read abundance is used to identify the most frequent 3′ end in each group. Second, corresponding MILI- and MIWI-bound pre-piRNA groups are paired and the distance is calculated between the most frequent 3′ end in MILI group and the most frequent 3′ end in MIWI group. If this distance is 0, the paired group falls into cohort 0; if the most frequent 3′ end in MILI group is 1 nt upstream of the most frequent 3′ end in MIWI group, the paired groups falls into cohort 1, etc.

**Figure S3, related to Figure 5.**
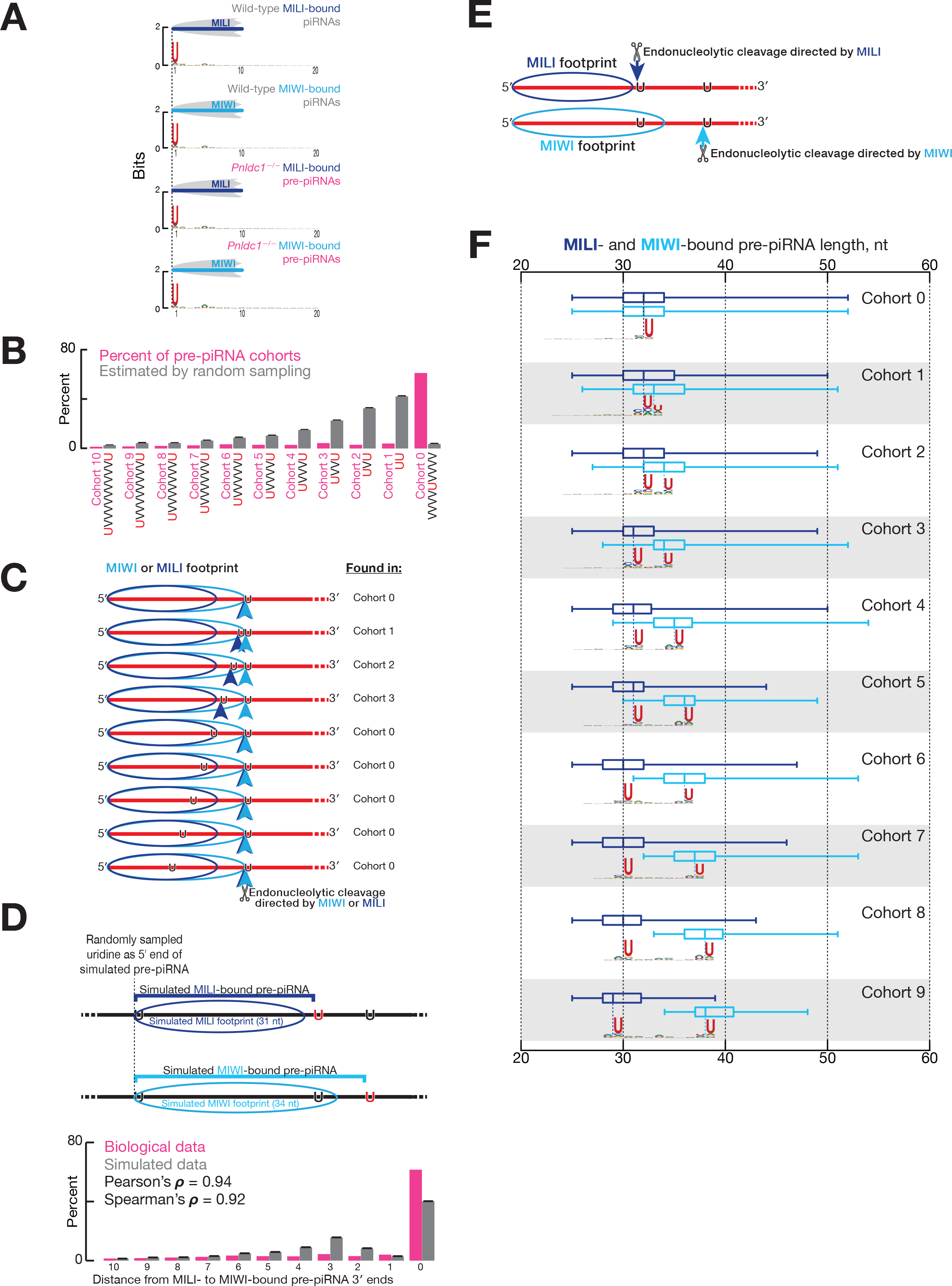
(A) Nucleotide bias for the 5′ section of MILI- or MIWI-bound piRNAs in wild-type and the same for pre-piRNAs in *Pnldc1^−/−^* primary spermatocytes. (B) Probability of a single uridine surrounded by non-uridine stretches (VVVVUVVVV), and probabilities of lengths of non-uridine nucleotide stretches between two nearest uridines (UU, UVU, UVVU, etc.) in pachytene piRNA loci estimated by random sampling of the data from pachytene piRNA cluster transcripts (grey bars). Mean ± standard deviation are presented for the randomly sampled data based on 1,000 iterations. Data for pre-piRNA cohorts (pink bars) are for a single biological sample from Figure 4C. (C) The first prediction of the model for pre-piRNA production in which MILI and MIWI proteins direct endonucleolytic cleavage by binding 5′ ends of pre-pre-piRNAs: protein footprints limit the availability of uridines for the pre-pre-piRNA cleaving endonuclease (vertical arrows). Because MILI footprint is smaller than that of MIWI, the endonuclease will have access to more upstream uridines if pre-pre-piRNA is bound by MILI compared to MIWI. MILI footprint will still place a 5′ limit on the upstream shift in the uridine availability windows for MILI and MIWI. (D) Test of the first prediction (Figure S3C) of the model for pre-piRNA production. Upper panel, strategy to create sets of simulated pre-piRNAs. Uridines randomly sampled in pachytene piRNA clusters were used as 5′ ends of simulated pre-piRNAs. The 3′ ends of the simulated pre-piRNAs were set immediately before the first uridine occurring >31 nt (simulated MILI footprint) or >34 nt (simulated MIWI footprint) downstream of the simulated pre-piRNA 5′ end. Lower panel, comparison of cohort sizes calculated from simulated and biological data. Biological data are from Figure 4C. Mean ± standard deviation are presented for the simulated data based on 1,000 iterations. (E) The second prediction of the model for pre-piRNA production in which MILI and MIWI proteins direct endonucleolytic cleavage by binding 5′ ends of pre-pre-piRNAs: MILI-bound pre-piRNAs present in cohorts ≥4 are paired with atypically long MIWI-bound pre-piRNAs because of the limitation on the minimal length of a MILI-bound pre-piRNA imposed by the MILI footprint. (F) Test of the second prediction (Figure S3E) of the model for pre-piRNA production. Length of the corresponding paired MILI- and MIWI-bound pre-piRNA groups in *Pnldc1^−/−^* primary spermatocytes in cohorts 0–9 (Figure 5). Data are from a single representative biological sample. Whiskers correspond to minimum and maximum values. Nucleotide bias of the genomic neighborhood around the 3′ ends of paired MILIand MIWI-bound pre-piRNA in each cohort is shown.

**Figure S4.**
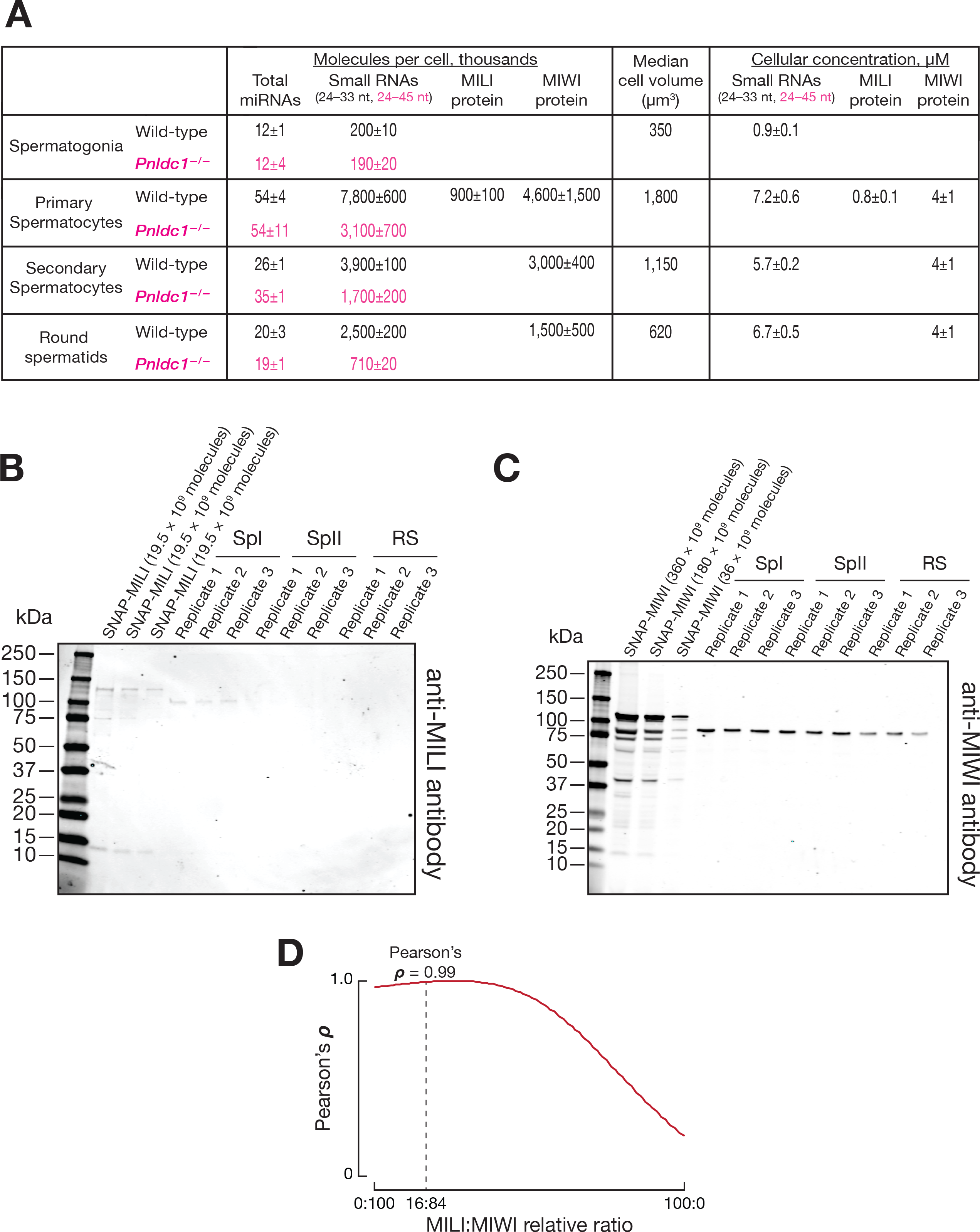
(A) Absolute abundance of miRNAs, small RNAs (24−33-nt for wild-type; 24−45-nt for *Pnldc1^−/−^*), and PIWI proteins at different stages of spermatogenesis in wild-type and *Pnldc1^−/−^* mice. Data are presented as the mean ± standard deviation from two biological replicates for small RNAs and from three biological replicates for MIWI proteins. Median cell volume was used to calculate cellular concentration of 24−45-nt long RNAs and PIWI proteins. (B, C) Abundance of MILI (B) and MIWI (C) in ~11,000 wild-type germ cells relative to the standards of SNAP-tagged PIWI proteins assessed by Western blotting. SpI, primary spermatocytes; SpII, secondary spermatocytes; RS, round spermatids. Data are from three biological samples. (D) Correlation between simulated and biological piRNA length distributions. Simulated profiles were created by combining MILI- and MIWI-bound piRNA length distributions at different ratios. Data are from a single representative biological sample.

**Figure S5, related to Figure 6.**
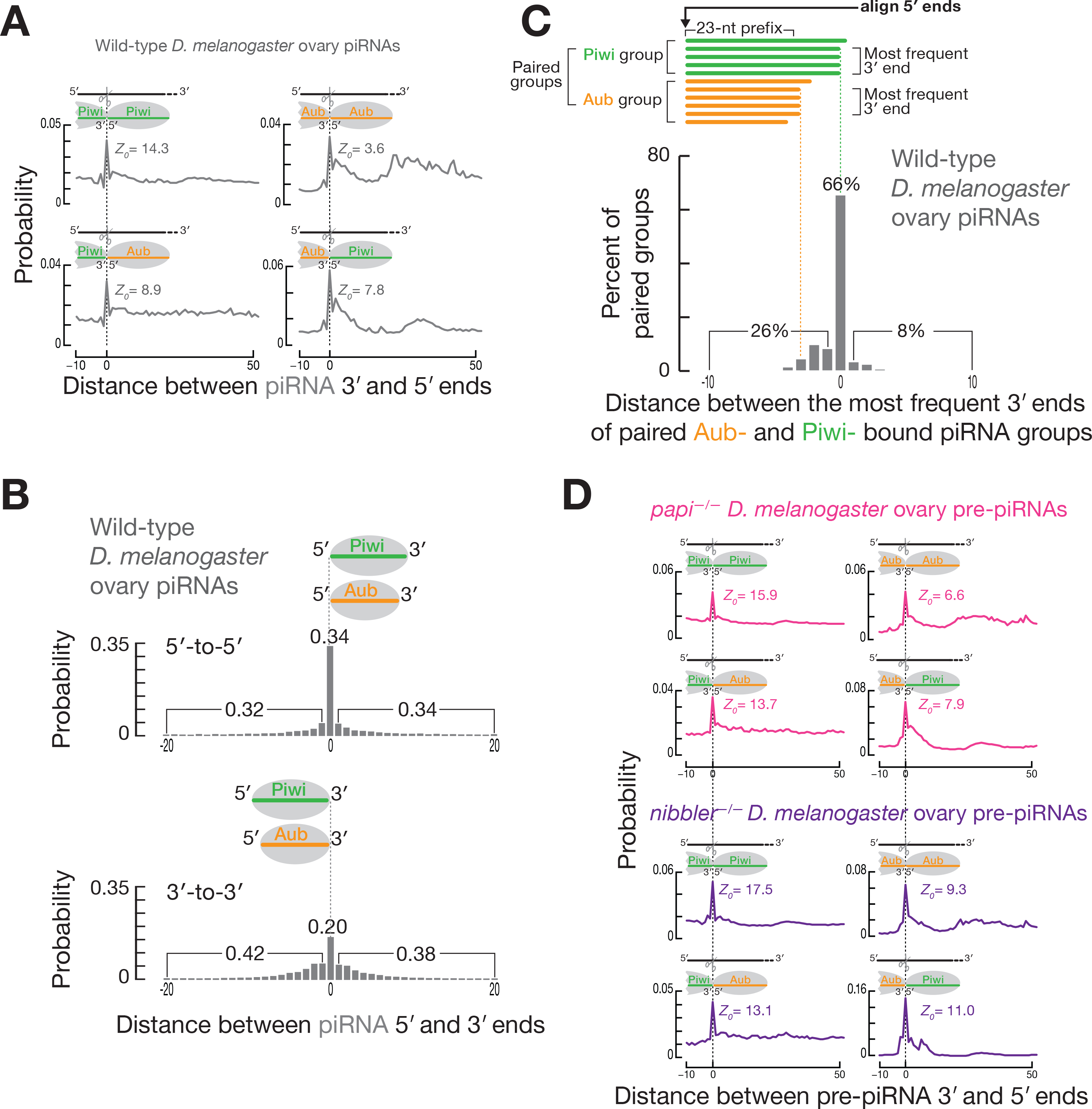
(A) Probability of distances from the 3′ ends of Piwi- or Aub-bound piRNAs to the 5′ ends of Piwi- or Aub-bound piRNAs on the same genomic strand in wild-type *D. melanogaster* ovary. Data are for all genome mapping piRNAs from a single representative biological sample. (B) Probability of distances from the 5′ ends of Piwi-bound piRNAs to the 5′ ends of Aub-bound piRNAs on the same genomic strand, and from the 3′ ends of Piwi-bound piRNAs to the 3′ ends of Aub-bound piRNAs on the same genomic strand in wild-type *D. melanogaster* ovary. Numbers indicate the total probability of 5′ or 3′ ends of Aub-bound piRNAs residing before, after or coinciding with the 5′ or 3′ ends of the Piwi-bound piRNAs. Data are for all genome mapping piRNAs from a single representative biological sample. (C) Distance between the most frequent 3′ end of the Piwi-bound piRNA group and the most frequent 3′ end of the corresponding paired Aub-bound piRNA group in wild-type *D. melanogaster* ovary. Data are for all unambiguously mapping piRNAs from a single representative biological sample. (D) Probability of distances from the 3′ ends of Piwi- or Aub-bound pre-piRNAs to the 5′ ends of Piwi- or Aub-bound pre-piRNAs on the same genomic strand for *papi^−/−^* and the same for *nibbler^−/−^ D. melanogaster* ovary. Data are for all genome mapping pre-piRNAs from a single representative biological sample.

**Figure S6, related to Figure 7.**
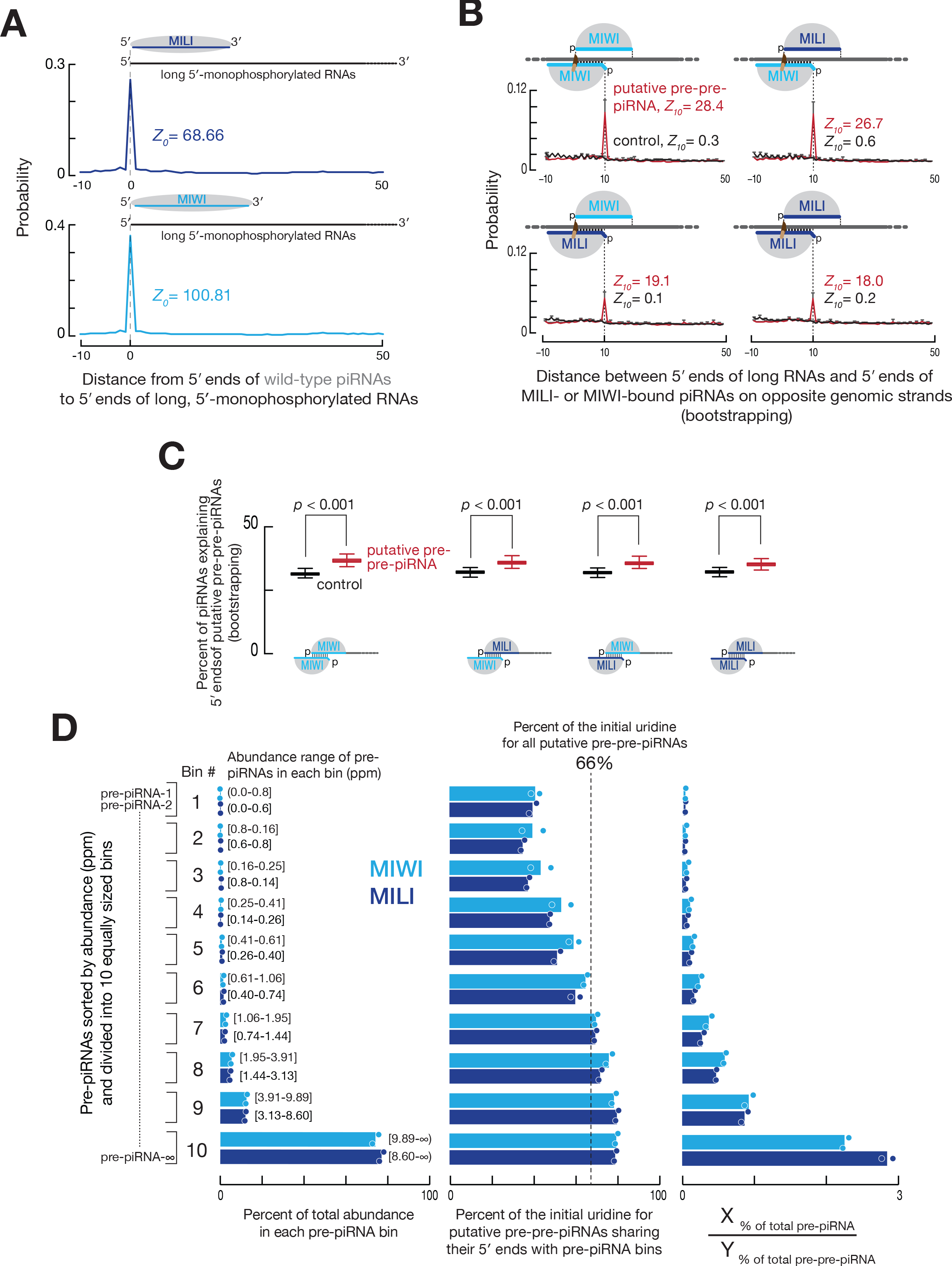
(A) Probability of distance from the 5′ ends of MILI- or MIWI-bound piRNAs to the 5′ ends of long 5′ monophosphorylated RNAs in wild-type primary spermatocytes. Data are for piRNAs deriving from pachytene piRNA loci from a single representative biological sample. (B) Probability of distances between the 5′ ends of long RNAs (putative pre-pre-piRNAs or control) and the 5′ ends of MILI- or MIWI-bound piRNAs on opposite genomic strands. Only nucleotides 2 to 10 of guide piRNAs (g2–g10) were required to be complementary to the target long RNAs. Data are for MILI- or MIWI-bound piRNAs derived from pachytene piRNA loci for a single biological sample and randomly resampled long RNAs derived from pachytene piRNA loci from the same biological sample. Mean ± standard deviation of 1,000 resampling iterations are presented. (C) Percent of piRNAs explaining the 5′ ends of either putative pre-pre-piRNAs or the control RNAs. Only nucleotides 2 to 10 of guide piRNAs (g2–g10) were required to be complementary to the target long RNAs. Data are for MILI- or MIWI-bound piRNAs derived from pachytene piRNA loci for a single biological sample and randomly resampled long RNAs derived from pachytene piRNA loci from the same biological sample. Data are from 1,000 resampling iterations. Whiskers correspond to minimum and maximum values. Wilcoxon rank-sum test was used to assess statistical significance. (D) Pre-piRNAs sorted by abundance and divided into 10 equally sized bins. At left, percent of total pre-piRNA abundance and the range of read abundance in each bin is shown. In the center, percent of the initial uridine among putative pre-pre-piRNAs sharing their 5′ ends with pre-piRNA bins. Percent of the initial uridine for all putative pre-pre-piRNAs is shown as the vertical dashed line. At right, ratio of the total pre-piRNA abundance to the total pre-pre-piRNA abundance in each bin.

**Figure S7.**
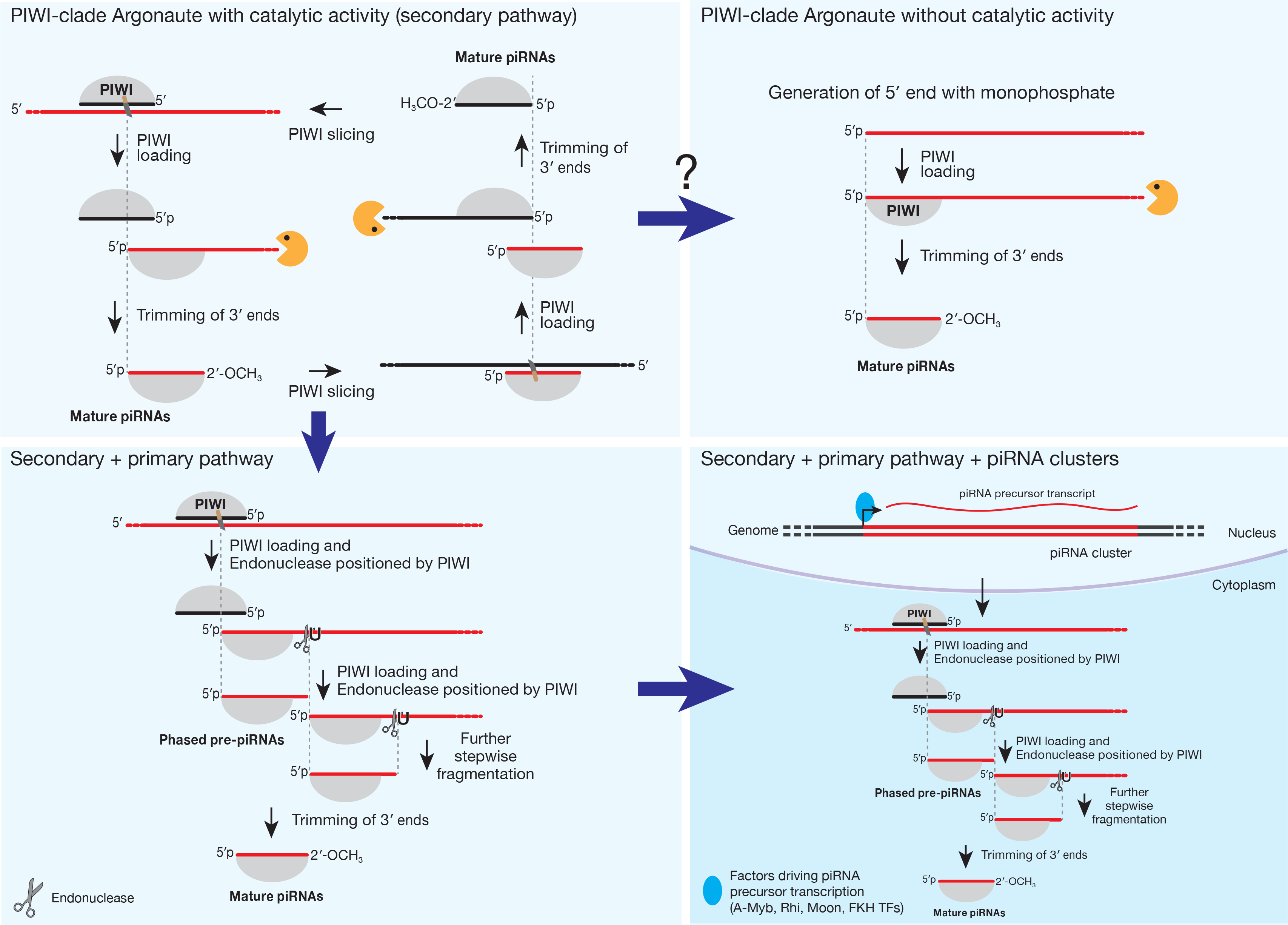
Proposed evolutionary trajectory of the piRNA pathway.

### STAR Methods

#### Mice

C57BL/6J mice (IMSR Cat# JAX:000664, RRID:IMSR_JAX:000664) were maintained and used according to the guidelines of the Institutional Animal Care and Use Committee of the University of Massachusetts Medical School. *Mili^−/−^* mice (IMSR Cat# RBRC09475, RRID:IMSR_RBRC09475) were a gift from Dr. Shinichiro Chuma (Kyoto University, Japan).

To create *Pnldc1^−/−^* mutants, three sgRNAs targeting sequences in exon 1, exon 2, and intron 2 of *Pnldc1* (5′-GUC CCA GGG CGC GCC GGA UG<u>C</u> <u>GG</u>)-3′, 5′-UGU CUC GGC CCC AAC AGA UC<u>A</u> <u>GG</u>)-3′, and 5′-UAA CUA AGG AGA CAC CGG UG<u>A</u> <u>GG</u>-3′, underlining denotes the PAM) were transcribed using T7 RNA Polymerase and purified by electrophoresis through a 10% polyacrylamide gel. Superovulated female C57BL/6J mice (7–8 weeks old) were mated to C57BL/6J stud males, and fertilized embryos were collected from oviducts. A mix of Cas9 mRNA (50 ng/µl, TriLink Biotechnologies, L-7206) and the three sgRNAs (each 20 ng/µl) were injected into the cytoplasm or pronucleus of fertilized eggs in M2 medium (Sigma, M7167). The injected zygotes were cultured in KSOM with amino acids at 37°C under 5% CO_2_ until the blastocyst stage (3.5 days). Thereafter, 15–25 blastocysts were transferred into the uterus of pseudopregnant ICR females at 2.5 dpc.

For genotyping, genomic DNA extracted from tail tissues was analyzed by PCR using primers 5′-TTC CCA GCA TGA GAA GAT CA-3′ and 5′-CCA CTC AGA TGG CAA GTC AA-3′. PCR products were Sanger-sequenced using the sequencing primer 5′-TGA CAC GTG CAC GAG CTT TA-3′. The male sterility phenotype reported previously (Ding et al., 2017; Zhang et al., 2017) was confirmed by the presence of epididymal sperm, testis weight, and histological assessment of the testes.

#### Isolation of Mouse Germ Cells by FACS

Testes were isolated, decapsulated, and rotated in 1× Gey′s Balanced Salt Solution (GBSS, Sigma, G9779) containing 0.4 mg/ml collagenase type 4 (Worthington LS004188) at 150 rpm for 15 min at 33°C. Seminiferous tubules were then washed twice with 1× GBSS and rotated in 1× GBSS with 0.5 mg/ml Trypsin and 1 µg/ml DNase I at 150 rpm for 15 min at 33°C. After incubation, the tubules were homogenized by pipetting with a Pasteur pipette for 3 min on ice, and fetal bovine serum (FBS; 7.5% f.c., v/v) was added to inactivate trypsin. The cell suspension was then strained through a pre-wetted 70 µm cell strainer and pelleted at 300× *g* for 10 min. The supernatant was removed, cells were resuspended in 1× GBSS containing 5% (v/v) FBS, 1 µg/ml DNase I, and 5 μgl/ml Hoechst 33342 (Thermo Fisher, 62249) and rotated at 150 rpm for 45 min at 33ºC. Propidium iodide (0.2 μg/ml, f.c.; Thermo Fisher, P3566) was added, and cells strained through a single pre-wetted 40 µm cell strainer. Cell sorting at the UMass Medical School FACS Core was performed as described previously (Bastos et al., 2005; Cole et al., 2014). Cell viability was assessed using live phase contrast microscopy. Purity check of sorted fractions was performed by immunostaining aliquots of cells.

For immunostaining, cells were incubated for 20 min in 25 mM sucrose and then fixed on a slide with 1% (w/v) paraformaldehyde solution containing 0.15% (v/v) Triton X-100 for 2 h at room temperature in a humidifying chamber. Slides were then sequentially washed for 10 min in (1) 1× PBS containing 0.4% (v/v) Photo-Flo 200 (Kodak, 1464510), (2) 1× PBS containing 0.1% (v/v) Triton X-100, (3) 1× PBS containing 0.3% (w/v) BSA, 1% (v/v) donkey serum (Sigma, D9663), and 0.05% (v/v) Triton X-100. After washing, slides were incubated with primary antibodies in 1× PBS containing 3% (w/v) BSA, 10% (v/v) donkey serum, and 0.5% (v/v) Triton X-100 in a humidifying chamber overnight at room temperature. Primary antibodies used in this study were rabbit polyclonal anti-SYCP3 (Abcam Cat# ab15093, RRID:AB_301639, 1:1000 dilution), and mouse monoclonal anti-γH2AX (Millipore Cat# 05-636, RRID:AB_309864, 1:1000 dilution). Slides were washed again as described above and incubated with secondary donkey anti-mouse IgG (H+L) Alexa Fluor 594 (Thermo Fisher Scientific Cat# A-21203, RRID:AB_2535789, 1:2000 dilution) or donkey anti-rabbit IgG (H+L) Alexa Fluor 488 (Thermo Fisher Scientific Cat# A-21206, RRID:AB_2535792, 1:2000 dilution) antibodies for 1 h in a humidifying chamber at room temperature. After incubation, slides were washed three times for 10 min in 1× PBS containing 0.4% (v/v) Photo-Flo 200 and once for 10 min in 0.4% (v/v) Photo-Flo 200. Finally, slides were dried, mounted with ProLong Gold Antifade Mountant with DAPI (Thermo Fisher, P36931), and covered with a cover slip.

#### SNAP-tagged MIWI Protein Standard

SNAP-tagged *Mus musculus* PIWIL1 (SNAP-MIWI) was produced in HEK293T cells from a lentiviral-transduced transgene (lentivirus backbone was a gift from Greiner Lab). Cells, washed twice with PBS, were homogenized in lysis buffer (30 mM HEPES-KOH pH 7.5, 100 mM potassium acetate, 3.5 mM magnesium acetate, 1 mM dithiothreitol, 20% (w/v) glycerol, 0.1% (v/v) Triton X-100, 1 mM 4-(2-Aminoethyl) benzenesulfonyl fluoride hydrochloride, 0.3 µM Aprotinin, 40 µM Bestatin, 10 µM E-64, 10 µM Leupeptin). Lysate was centrifuged at 20,000 × *g* for 30 min, and the supernatant aliquoted and stored at −80°C. SNAP-MIWI was labeled with SNAP substrate SNAP-Surface 549 (NEB, S9112) and resolved by electrophoresis through a 4–20% gradient SDS-polyacrylamide gel (Bio-Rad Laboratories, 5671085). The concentration of full-length SNAP-MIWI was determined by comparison to a standard curve of purified, SNAP-Surface 549 labeled SNAP protein (Typhoon FLA 7000; GE Lifesciences). The concentration of purified SNAP protein (gift from Moore Lab) was determined before labeling by BCA assay and by measuring its absorbance at 280 nm (ε = 20970 M^-1^ cm^-1^ in water assuming all Cys residues are reduced). SNAP-MIWI of known concentration was then used to estimate the number of MIWI molecules present in FACS-purified germ cells from mouse testes by quantitative immunoblotting using anti-MIWI antibody.

#### Western Blotting

Cells were homogenized in lysis buffer (20 mM Tris-HCl pH 7.5, 2.5 mM MgCl_2_, 200 mM NaCl, 0.05% (v/v) NP-40, 0.1 mM EDTA, 1 mM 4-(2-Aminoethyl) benzenesulfonyl fluoride hydrochloride, 0.3 µM Aprotinin, 40 µM Bestatin, 10 µM E-64, 10 µM Leupeptin) and centrifuged at 20,000 × *g* for 20 min at 4°C. The supernatant was moved to a new tube, an equal volume of loading dye (120 mM Tris-Cl, pH 6.8, 4% (w/v) SDS, 20% (v/v) glycerol, 2.5% (v/v) 2-Mercaptoethanol, 0.2% (w/v) bromophenol blue) was added, the sample incubated at 90°C for 5 min and resolved through a 4–20% gradient polyacrylamide/SDS gel electrophoresis (Bio-Rad Laboratories, 5671085). After electrophoresis, proteins were transferred to PVDF membrane (Millipore, IPVH00010), the membrane blocked in Blocking Buffer (Rockland Immunochemicals, MB-070) at room temperature for 2 h and then incubated overnight at 4°C in Blocking Buffer containing primary antibody (anti-PIWIL2/MILI, Abcam Cat# ab36764, RRID:AB_777284, 1:1000 dilution; anti-PIWIL1/MIWI, Abcam Cat# ab12337, RRID:AB_470241, 1:1000 dilution). Next, the membrane was washed three times with Blocking Buffer at room temperature for 30 min and incubated for 2 h at room temperature with donkey anti-rabbit IRDye 680RD secondary antibody (LI-COR Biosciences Cat# 926-68073, RRID:AB_1095444, diluted 1:20,000) in Blocking Buffer. Then the membrane was washed three times with Blocking Buffer at room temperature for 30 min and the signal was detected using Odyssey Infrared Imaging System. Data was obtained for two independent biological replicates.

#### Small RNA Immunoprecipitation

Mouse total testis or sorted germ cells were homogenized with lysis buffer (20 mM TrisHCl, pH 7.5, 2.5 mM MgCl_2_, 200 mM NaCl, 0.05% (v/v) NP-40, 0.1 mM EDTA, 1 mM 4-(2-Aminoethyl) benzenesulfonyl fluoride hydrochloride, 0.3 µM Aprotinin, 40 µM Bestatin, 10 µM E-64, 10 µM Leupeptin) and then centrifuged at 20,000 × *g* for 20 min at 4°C, retaining the supernatant. Anti-MIWI (Wako, Cat# 017-23451, RRID:AB_2721829, ~5 µg; Nishibu, 2012) or anti-MILI (Abcam Cat# ab36764, RRID:AB_777284, ~5 µg) antibodies were incubated with rotation with 30 µl of Protein G Dynabeads (Thermo Fisher, 10003D) in 1× PBS containing 0.02% (v/v) Tween 20 (PBST) at 4°C for 1 h. The bead-antibody complex was washed with PBST. Freshly prepared testis or cell lysate were added to the bead-antibody complex and incubated with rotation at 4°C overnight. The next day, the beads were washed once with lysis buffer and three times with 0.1 M Trisodium Citrate. After washing, RNA was purified with Trizol reagent (Thermo Fisher, 15596026) and used for small RNA library preparation. Each experiment was conducted for two independent biological replicates. The specificity of the commercial anti-MILI antibody was confirmed by immunoprecipitation from lysate of *Mili^−/−^* whole testis (data not shown).

#### Small RNA-seq Library Preparation and Analysis of Small RNA Data Sets

Total RNA from sorted mouse germ cells was extracted using mirVana miRNA isolation kit (Thermo Fisher, AM1560). Small RNA libraries were constructed as described (Han et al., 2015) with several modifications. Briefly, before library preparation, a set of 18 synthetic RNA oligonucleotides was added to each RNA sample to enable absolute quantification of small RNAs (Table S2A). To reduce ligation bias, a 3′ adaptor with three random nucleotides at its 5′ end was used (5′-rApp NNN TGG AAT TCT CGG GTG CCA AGG /ddC/-3′). After 3′ adaptor ligation, RNA was purified by 15% urea polyacrylamide gel electrophoresis (PAGE), selecting for 15–55 nt small RNAs (i.e., 40–80 nt with 3′ adaptor). Small RNA-seq libraries for two independent biological replicates were sequenced using a NextSeq 500 (Illumina) to obtain 75 nt, single-end reads.

The sequence of the 3′ adapter, including the three random nucleotides, was removed from raw reads, which were further filtered by requiring their Phred quality score to be ≥20 for all nucleotides. Sequences of synthetic spike-in oligonucleotides were identified allowing no mismatches (Table S2A). Absolute quantity of small RNAs per library was calculated based on the read abundance of 2′-*O*-methylated synthetic spike-in oligonucleotides (Table S2B). Reads not fully matching the genome were analyzed using the Tailor pipeline (Chou et al., 2015) to account for non-templated tailing of small RNAs.

*Drosophila melanogaster* Piwi- and Aub-bound small RNA data sets used in this study are listed in Table S3A. Distances between small RNAs were calculated using all possible alignments of either in all genome-matching reads (mouse and fly data sets) or matching only to annotated pachytene piRNA loci (mouse data sets only; Li et al., 2013), taking into account the number of times a small RNA sequence occurs in the library divided by the number of locations where this small RNA maps in the genome (i.e., multi-mapping reads were apportioned). *Z_0_* scores for phasing were calculated as described (Han et al., 2015) using distances from –10 to –1 and from 1 to 50 as background. Sequence motif charts for genomic neighborhoods around 5′ and 3′ ends of small RNAs were generated with motifStack (Ou et al., 2018) using alignments in prepachytene or pachytene piRNA loci only (Li et al., 2013) and apportioning reads.

Grouping of piRNAs or pre-piRNAs (≥1 ppm) with the same 5′, 23-nt or 25-nt prefix was done for all unambiguously mapping piRNAs (mouse and fly data sets) or for those mapping to the pachytene piRNA loci only (mouse data sets only; Li et al., 2013). The most frequent 3′ end in each group was identified based on the number of times 3′ ends are found in the library. Pairing MILI- and MIWI-bound small RNA groups was done based on their 5′, 25-nt prefix. Pairing Aub- and Piwi-bound small RNA groups was done based on their 5′, 23-nt prefix.

Distance probability analyses of previously published datasets from 34 animal species (Table S3B; Grimson et al., 2008; Friedländer et al., 2009; Kamminga et al., 2010; Song et al., 2012; Li et al., 2013; Juliano et al., 2014; Hirano et al., 2014; Moran et al., 2014; Xu et al., 2014; Han et al., 2015; Williams et al., 2015; Sarkies et al., 2015; Roovers et al., 2015; Rosenkranz et al., 2015; Toombs et al., 2017; Lewis et al., 2017; Fu et al., 2018) were done for ≥24-nt sequencing reads without taking into account their abundance. Smoothing of the data points [20-200] with non-parametric regression (LOWESS) was conducted in R without robustifying iterations and the span set at 0.1. Ping-pong *Z_10_* score was calculated as described (Zhang et al., 2011) for ≥24-nt sequencing reads without taking into account their abundance using distances from 0 to 9 and from 11 to 20 as background.

#### RNA-seq Library Preparation and Analysis

Total RNA from sorted germ cells was extracted using mirVana miRNA isolation kit (Thermo Fisher, AM1560) and used for library preparation as described previously (Zhang et al., 2012) with several modifications (Fu et al., 2018). Briefly, before library preparation, 1 µl of 1:100 dilution of ERCC spike-in mix 1 (Thermo Fisher, 4456740, LOT00418382) was added to 0.5–1 µg total RNA to enable absolute quantification of mRNA. For ribosomal RNA depletion, RNA was hybridized in 10 µl to a pool of 186 rRNA antisense oligos (0.05 µM each; Morlan et al., 2012; Adiconis et al., 2013) in 10 mM Tris-Cl (pH 7.4), 20 mM NaCl, heating the mixture to 95°C, cooling it at –0.1°C/sec to 22°C, and incubating at 22°C for 5 min. RNase H (10U; Lucigen, H39500) was added and the mixture incubated at 45°C for 30 min in 20 µl containing 50 mM Tris-Cl (pH 7.4), 100 mM NaCl, and 20 mM MgCl_2_. RNA was further treated with 4U DNase (Thermo Fisher, AM2238) in 50 µl at 37°C for 20 min. After DNase treatment, RNA was purified using RNA Clean & Concentrator-5 (Zymo Research, R1016). RNA-seq libraries for three independent biological replicates were sequenced using a NextSeq 500 (Illumina) to obtain 75 + 75 nt, paired-end reads.

RNA-seq analysis was performed with piPipes (Han et al., 2014). Briefly, RNAs were first aligned to ribosomal RNA sequences using Bowtie2 (v2.2.0; Langmead and Salzberg, 2012). Unaligned reads were then mapped using STAR to mouse genome mm10 (v2.3.1; Dobin et al., 2013). Sequencing depth and gene quantification was calculated with Cufflinks (v2.1.1; Trapnell et al., 2010). Differential expression analysis for piRNA precursor transcripts (Li et al., 2013) was performed using DESeq2 (v1.18.1; Love et al., 2014). In parallel, raw reads were aligned to an index of ERCC spike-in transcripts (Thermo Fisher, 4456740, LOT00418382) using Bowtie (v1.0.0; Langmead et al., 2009) to enable assessment of the absolute quantity of transcripts in each library.

#### Cloning and Sequencing of 5′ monophosphorylated Long RNAs and Analysis of Sequencing Data

Total RNA from sorted mouse germ cells was extracted using mirVana miRNA isolation kit (Thermo Fisher, AM1560) and used to prepare a library of 5′ monophosphorylated long RNAs as described (Wang et al., 2014). Libraries for two independent biological replicates were sequenced using a NextSeq 500 (Illumina) to obtain 75 + 75 nt, paired-end reads. Bioinformatics analysis was performed with piPipes (Han et al., 2014). Briefly, RNAs were first aligned to ribosomal RNA (rRNA) sequences using Bowtie2 (v2.2.0; Langmead and Salzberg, 2012). Unaligned reads were then mapped using STAR to mouse genome mm10 (v2.3.1; Dobin et al., 2013) and alignments with soft clipping of ends were removed with SAMtools (v1.0.0; Li et al., 2009; Li, 2011). The number of overlaps between small RNAs and the 5′ ends of long RNAs on opposite strands was calculated with read apportioning, analyzing only sequences mapping to the pachytene piRNA loci (Li et al., 2013). Ping-pong *Z_10_* score was calculated as described (Zhang et al., 2011) using distances from –10 to 9 and from 11 to 50 as background. Bootstrapping was performed with 1,000 iterations for randomly resampled sets of 5′ monophosphorylated long RNAs. The size of the resampled sets (100,000 species) was based on the median size of sets of 5′ monophosphorylated long RNAs.

## SUPPLEMENTAL ITEM TITLES

**Table S1. Relative and Absolute Steady-State Levels of Transcripts.**

Data are for all annotated genes in spermatogonia, primary spermatocytes, secondary spermatocytes and round spermatids of wild-type and *Pnldc1^−/−^* mice. Relative data are presented in fpkm (Fragments Per Kilobase of transcript per Million mapped reads), absolute data are presented in molecules per cell.

**Table S2. Synthetic Spike-in RNA Oligonucleotides.**

(A) Sequences of synthetic spike-in RNA oligonucleotides used to measure small RNA abundance.

(B) Amount of synthetic spike-in RNA oligonucleotides and the total number of cells used for the preparation of each small RNA library.

**Table S3. Data Sets Used in This Study**

(A) *Drosophila melanogaster* Piwi- and Aub-bound small RNA data sets used in this study.

(B) Small RNA data sets from 34 animal species used for analyses presented in Figure 2.

